# PfCERLI1, a conserved rhoptry associated protein essential for invasion by *Plasmodium falciparum* merozoites

**DOI:** 10.1101/646299

**Authors:** Benjamin Liffner, Sonja Frölich, Gary K. Heinemann, Boyin Liu, Matthew W. A. Dixon, Tim-Wolf Gilberger, Danny W. Wilson

## Abstract

The disease-causing blood stage of the *Plasmodium falciparum* lifecycle begins with invasion of human erythrocytes by merozoites. Many vaccine candidates with key roles in binding to the erythrocyte surface and entry are secreted from the large bulb-like rhoptry organelles at the apical tip of the merozoite. Here we identify an essential role for the conserved protein *P. falciparum* <u>C</u>ytosolically <u>E</u>xposed <u>R</u>hoptry <u>L</u>eaflet <u>I</u>nteracting protein 1 (PfCERLI1) in rhoptry function. We show that PfCERLI1 localises to the cytosolic face of the rhoptry bulb membrane and knockdown of PfCERLI1 inhibits merozoite invasion. While schizogony and merozoite organelle biogenesis appear normal, biochemical techniques and semi-quantitative super-resolution microscopy show that PfCERLI1 knockdown prevents secretion of key rhoptry antigens that coordinate merozoite invasion. PfCERLI1 is the first rhoptry associated protein identified to have a direct role in function of this essential malaria invasion organelle which has broader implications for understanding apicomplexan invasion biology.

## Introduction

Malaria, caused by infection with *Plasmodium* spp. parasites, results in >400,000 deaths annually with *Plasmodium falciparum* being responsible for the majority of malaria mortality^1^. *Plasmodium* parasites have a complex lifecycle with human infection beginning with transmission of the liver-cell invading sporozoite from a female Anopheline mosquito to the human host. Thousands of daughter merozoites develop in the maturing liver-stage parasite and are then released into the blood stream where they invade human erythrocytes. In the case of *P. falciparum*, the blood stage parasite grows and multiplies over the next 48 hours, forming 16-32 daughter merozoites that rupture out of the late schizont stage parasite and infect new erythrocytes^2^. This asexual blood stage of the malaria parasite lifecycle causes all symptoms of clinical disease.

Erythrocyte invasion, the first step in blood stage infection, is an essential and highly co-ordinated process that begins with reversible attachment of the merozoite to the erythrocyte surface^3^. The merozoite then reorientates such that the apical tip, containing specialised invasion organelles known as the rhoptries and micronemes, comes in contact with the erythrocyte membrane. Invasion ligands and adhesins are released from the micronemes and rhoptries prior to formation of an irreversible tight-junction that binds the merozoite to the erythrocyte surface^4^. The merozoite then engages an actin-myosin based motor complex, anchored in an inner membrane complex (IMC) that lies under the merozoite plasma membrane, to pull the erythrocyte membrane around itself; thus completing invasion^2,5,6^.

Merozoite invasion is a key stage of the parasite lifecycle with the potential to be targeted by therapeutic interventions that would prevent blood-stage parasite replication and disease^3,5,7–10^. It is thought, in *Plasmodium falciparum*, that the contents of micronemes are released first and are responsible for coordinating initial attachment events^11^. By contrast, the contents of the dense granules are typically involved in host-cell remodelling following the completion of invasion^12–14^. The rhoptries, which are the largest of the invasion organelles, adopt a dual club-shape with bulbs further towards the basal side of the merozoite and necks at the very apical tip^15^. During invasion, the contents of the rhoptry neck are released first and initiate early interactions with host erythrocyte receptors^11,16^. Rhoptry bulb contents are secreted later in the invasion process and are typically involved in establishment of the parasitophorous vacuole^17,18^.

It is currently unclear what differentiates the rhoptry bulb from the neck, as no membrane separates them. Because the neck and bulb remain differentiated even after plasma membrane fusion, it has been hypothesised that the structure of the rhoptry neck and bulb may be supported by a scaffold but the components of such a scaffold are yet to be elucidated^19^. Additionally, it is thought that there is at least partial sub-compartmentalisation of proteins within the rhoptry bulb and neck and that this may be related to the timing of protein release^20^. Secretion of rhoptry contents may be partially coordinated through changes in the organelle’s structure during invasion. The membranes at the tip of the rhoptry neck are fused with the parasite membrane early in invasion and the two rhoptries themselves also fuse beginning at the neck^19^. As invasion proceeds the rhoptries fuse together fully into a single rhoptry (neck and bulb) at the point of erythrocyte entry. However, the precise mechanisms leading to expulsion of luminal rhoptry contents are yet to be elucidated.

To date, approximately 30 proteins have been linked to the rhoptry^15^. Despite the rhoptries being membrane-bound organelles with dynamic function during secretion of invasion ligands, there remains only a single protein reported to localise to either the inner (Rhoptry associated membrane antigen) or outer (Armadillo repeats only) surfaces of the *P. falciparum* rhoptry membrane. Therefore, the drivers of rhoptry secretion and compartmentalisation are unknown. Here we describe a highly conserved *P. falciparum* protein, Pf3D7_0210600 (henceforth named *Plasmodium falciparum* <u>C</u>ytosolically <u>E</u>xposed <u>R</u>hoptry <u>L</u>eaflet <u>I</u>nteracting protein (PfCERLI1)), that lies on the cytosolic face of the rhoptry bulb and has an essential role in rhoptry secretion and merozoite invasion.

## Results

### PfCERLI1 is highly conserved among *Plasmodium* spp. and is essential for blood-stage growth

PfCERLI1 is a protein of 446 amino acids that is predicted to contain a signal peptide but no transmembrane domains or glycophosphatidylinositol (GPI) anchor that would clearly mark it as a membrane protein (Fig. 1a). Previous studies have shown that PfCERLI1 may be enriched by acyl biotin exchange, suggesting that it may be palmitoylated^21^. The RNA expression profile and predicted signal peptide of PfCERLI1 has led to speculation that it has a role in merozoite invasion that could be targeted by vaccines^22,23^. PfCERLI1 is highly conserved among *Plasmodium* spp., with >90% identity between *P. falciparum*, *P. reichenowi* and *P. gaboni* orthologues, >75% amino acid identity evident across human infecting species (including the zoonotic *P. knowlesi*) and >70% when compared to distantly related species (Fig. 1b). To determine whether this high inter-species conservation was due to a function essential for blood-stage survival, we attempted to disrupt the *Pfcerli1* gene using the selection linked integration-targeted gene disruption system (SLI-TGD) (Fig. 1c)^24^. Our attempts to disrupt *Pfcerli1* were unsuccessful, supporting the results of a random mutagenesis study^25^, that suggest PfCERLI1 is essential for *P. falciparum* blood-stage growth.

**Fig 1.**
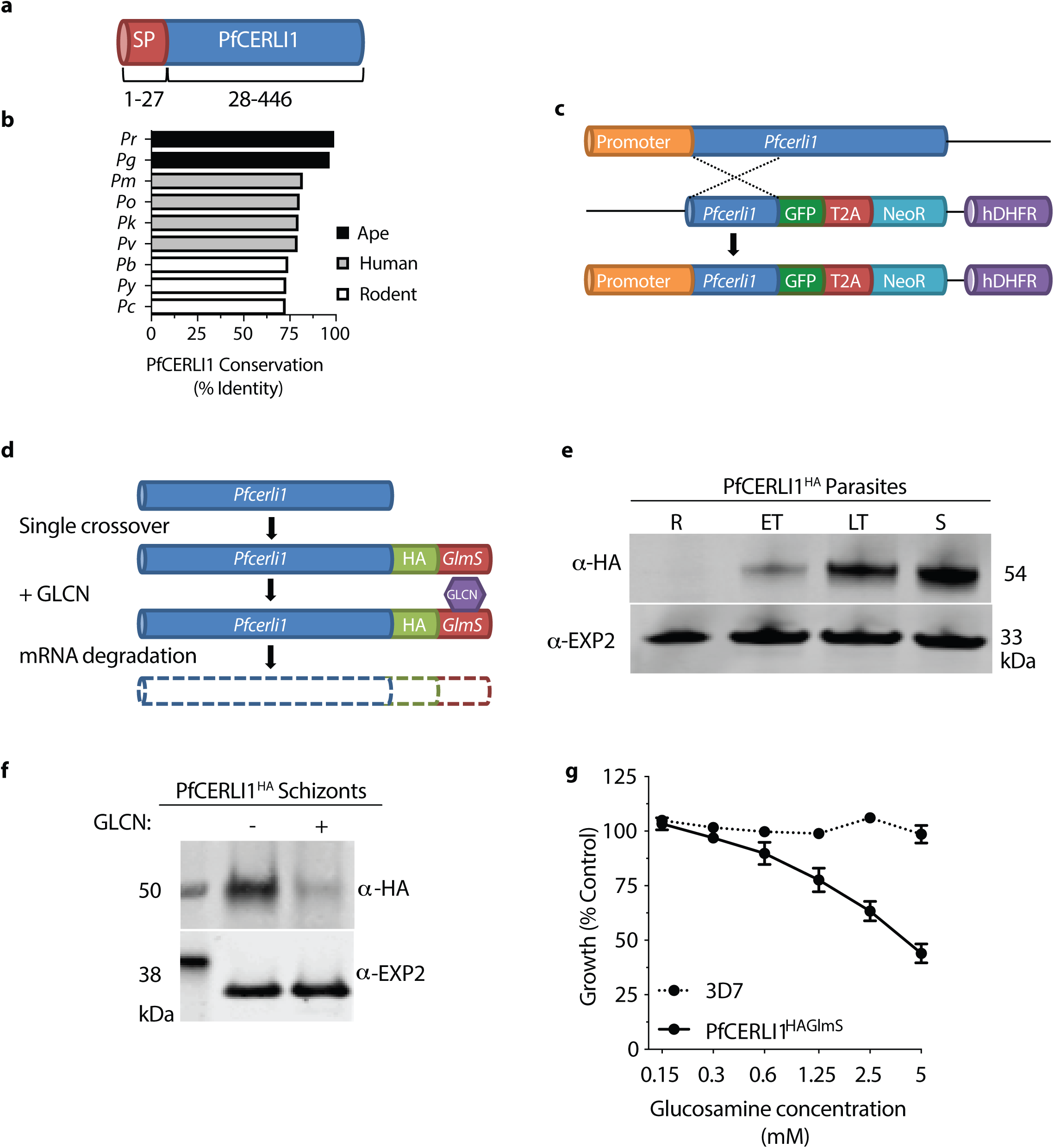
Phylogeny of PfCERLI1 (Pf3D7_0210600) and development of genetic tools to investigate function. **(a)** The annotated PfCERLI1 protein is 446 amino acids in length and is predicted to contain a signal peptide. **(b)** The amino acid sequence of PfCERLI1 was compared against predicted *Plasmodium* spp. orthologues by multiple pairwise alignments. *Pr* = *P. reichenowi*, *Pg* = *P. gaboni*, *Pm* = *P. malariae*, *Po* = *P. ovale*, *Pk* = *P. knowlesi*, *Pv* = *P. vivax*, *Pb* = *P. berghei, Py* = *P. yoelii*, *Pc* = *P. chabaudi*. **(c)** Schematic representation of the selection linked integration targeted gene disruption (SLI-TGD) system used to attempt knock-outs of *Pfcerli1*. A plasmid containing a *Pfcerli1* flanking region, green fluorescent protein (GFP) a T2A skip peptide and neomycin resistance cassette (NeoR) was transfected into wildtype 3D7 parasites. Single cross-over integration (knock-out) is expected to be driven through selection of neomycin resistance since this can only occur when NeoR expression is driven by the endogenous *Pfcerli1* promoter. Plasmid uptake was selected using the human dihydrofolate reductase (hDHFR) cassette, which confers resistance to the drug WR99210. **(d)** Schematic representation of the HA *GlmS* riboswitch system used to study PfCERLI1. A plasmid vector, containing a 3’ flank of the *Pfcerli1* sequence with a haemagglutinin (HA) tag and under the control of a *GlmS* ribozyme was transfected into wildtype parasites by 3’ single crossover recombination. Glucosamine (GLCN) binds to the *GlmS* ribozyme, specifically promoting the degradation of PfCERLI1 mRNA. **(e)** Western blot of ring (R), early trophozoite (ET), late trophozoite (LT) or schizont (S) stage PfCERLI1^HAGlmS^ lysates that were probed with anti-HA (PfCERLI1) or anti-EXP2 (loading control) antibodies. **(f)** Western blot of schizont stage PfCERLI1^HAGlmS^ lysates either in the presence (+) or absence (-) of 2.5 mM GLCN, which was then probed with anti-HA (PfCERLI1) and anti-EXP2 (loading control) antibodies, showing effective knockdown of PfCERLI1 in the presence of GLCN. **(g)** Synchronous PfCERLI1^HAGlmS^ or 3D7 trophozoite-stage parasites were treated with increasing concentrations of GLCN for 48 hours, with the number of trophozoites the following cycle measured to determine knockdown-mediated growth inhibition (parasite growth expressed as a % of non-inhibitory media controls, n=3, error bars = standard error of the mean (SEM)). X-axis presented as a log 2 scale for viewing purposes.

### PfCERLI1 is expressed in late stage schizonts and its loss is inhibitory to blood-stage growth

In order to analyse protein expression levels and function of PfCERLI1 during blood stage development, we made transgenic *P. falciparum* parasites that express a C-terminal haemagglutinin tagged (HA) PfCERLI1 with control of protein expression achieved through a glucosamine (GLCN) inducible *GlmS* ribozyme knockdown system (PfCERLI1^HAGlmS^)^26,27^ (Fig. 1d). Using lysates prepared from synchronised blood-stage parasite cultures (harvested at rings, early trophozoites, late trophozoites and schizonts) probed with an anti-HA antibody, we determined that the HA-tagged PfCERLI1 protein was most highly expressed in late schizont stages (Fig. 1e), concordant with previously published transcriptomic data^28^.

To validate that the integrated *GlmS* ribozyme could control PfCERLI1 protein expression, we treated PfCERLI1^HAGlmS^ parasites with 2.5 mM GLCN and quantified changes in HA-tagged protein levels using Western blot. Treatment of synchronous parasites with 2.5 mM GLCN for ~44 hours from early ring stage led to a >80% reduction in PfCERLI1 expression, whereas expression of the loading control EXP2 was not affected (Fig. 1f; Supplementary Fig. 1). To determine whether loss of PfCERLI1 protein expression affected parasite growth, early ring stage PfCERLI1^HAGlmS^ parasites were treated with increasing concentrations (0.125 to 5 mM) of GLCN for 48 hours and parasite growth assessed the following cycle. Treatment of PfCERLI1^HAGlmS^ with increasing concentrations of GLCN for 48 hours caused dose-dependent growth inhibition, with 5 mM GLCN leading to an approximately 55% reduction in parasite growth (Fig. 1g). By contrast, GLCN treatment did not lead to significant growth inhibition in control 3D7 parasites. Using a concentration of 2.5 mM, which has minimal non-specific growth inhibitory activity, PfCERLI1^HAGlmS^ blood-stage parasite growth was reduced by 40% after 48 hours treatment. Combined with PfCERLI1 being refractory to gene deletion and its high level of conservation, these data show that PfCERLI1 has an important role in asexual stage parasite growth and the stage of protein expression suggested that it could be involved in merozoite invasion of new RBCs.

### PfCERLI1 has an important role in merozoite invasion

In order to assess whether PfCERLI1 has a role in merozoite invasion, we co-transfected the PfCERLI1^HAGlmS^ parasites with a cytosolic green fluorescent protein (GFP) reporter plasmid (PfCERLI1^HAGlmS/GFP^) that allowed accurate quantitation of new merozoite invasion events by flow cytometry^29,30^. Since PfCERLI1 was expressed only at schizont stages, we limited the amount of time PfCERLI1^HAGlms/GFP^ parasites were exposed to increasing concentrations of GLCN (0.125 to 5 mM) by only treating from early trophozoite stages for 24 hours before assessing parasite rupture and invasion. There was a dose-dependent inhibition of merozoite invasion (Fig. 2a) and this inhibition was of a similar magnitude to levels of growth inhibition recorded with longer term treatment, strongly suggesting that knockdown of PfCERLI1 restricts parasite growth by preventing invasion.

**Fig 2.**
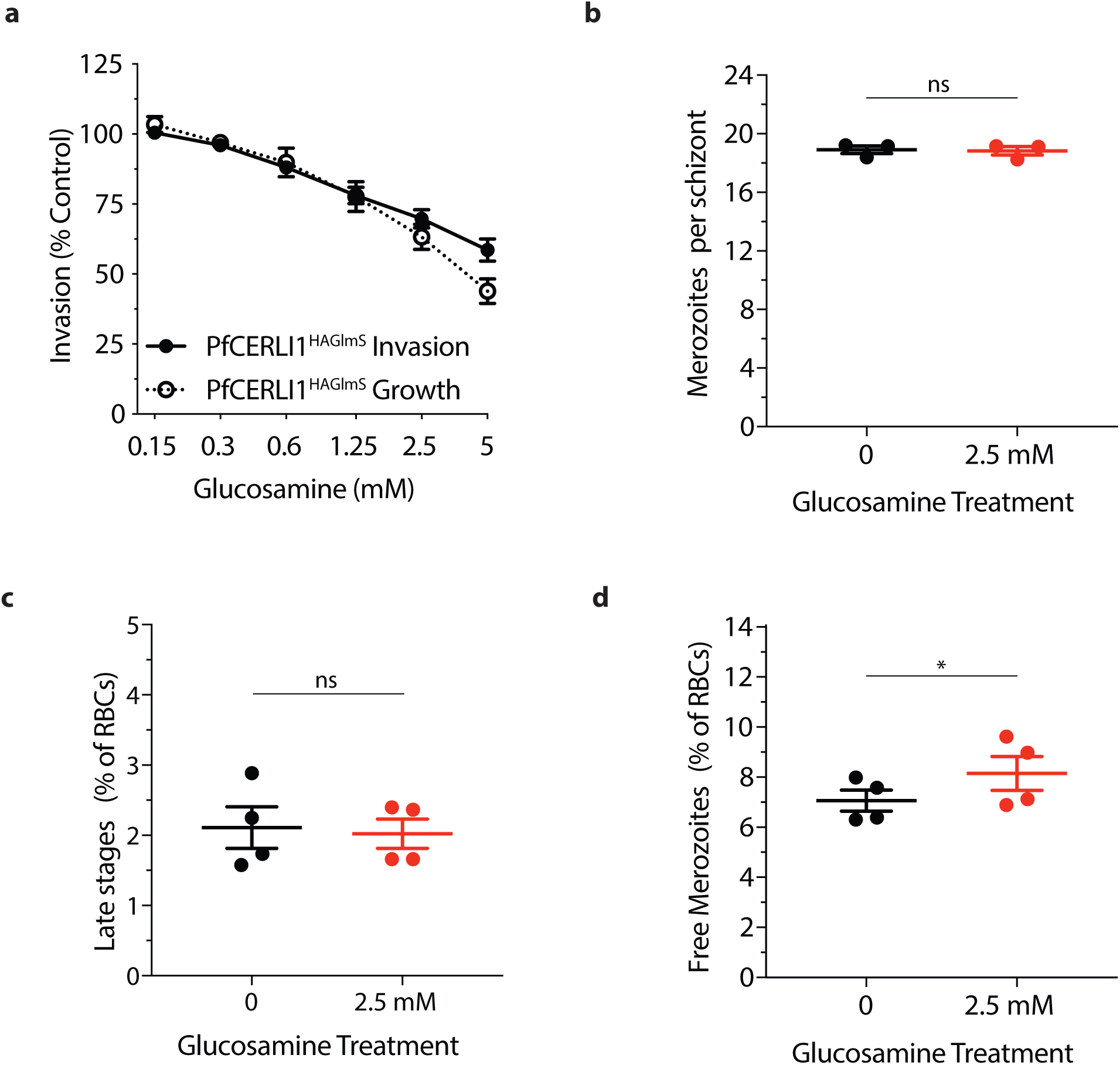
PfCERLI1 knockdown does not inhibit merozoite development but does prevent merozoite invasion. **(a)** Flow cytometric detection of GFP-expressing PfCERLI1^HAGlmS/GFP^ ring stage parasites after merozoite invasion indicated a direct inhibition of merozoite invasion with protein knockdown (results presented as a % of media control, n=4). PfCERLI1^HAGlmS^ Growth is replicated from Figure 1g for direct comparison between growth and invasion inhibition. X-axis presented as log 2 scale for viewing purposes. **(b)** Mean number of fully segmented merozoites per schizont was determined by microscopy analysis of Giemsa-stained thin blood smears. Smears were blinded and counted in three independent treatments with each data point representing the mean number of merozoites per schizont pooled from 20 schizonts. The number of unruptured trophozoites **(c)** and free merozoites **(d)** after GLCN treatment was assessed using flow cytometry, with results presented as % of total erythrocytes. Each data point represents triplicate wells from four independent experiments (ns = p>0.05, * = p<0.05 by Mann-Whitney or paired t-test, n = 4, error bars = SEM).

In order to define if knockdown of PfCERLI1 was inhibiting invasion by interfering with merozoite production, the number of merozoites that fully developed per E64 treated PfCERLI1^HAGlmS/GFP^ schizont was counted using light-microscopy of Giemsa stained thin smears. There was no difference in the number and stage of merozoite development evident between GLCN treated and untreated PfCERLI1^HAGlmS/GFP^ parasites (Untreated 18.9 merozoites per schizont; treated 18.8 merozoites per schizont, p=0.5; Fig. 2b). Furthermore, flow cytometry analysis also indicated that the number of unruptured schizonts was not altered with GLCN treatment (p=0.6) (Fig. 2c). When the number of free merozoites between GLCN and non-treated cultures was compared, there was significantly more free merozoites in the GLCN treated cultures (p<0.05) (Fig. 2d). When the number of lost invasion events (lost ring stage parasitaemia) was subtracted from the number of free merozoites, there was no difference between GLCN treated and untreated parasites (p>0.99, Supplementary Fig. 2a). These data indicate that knockdown of PfCERLI1 is associated with a loss of merozoite invasion prior to formation of the tight-junction, leading to an increase in the merozoite population free-floating in the culture medium. We also assessed whether knockdown of PfCERLI1 protein expression could lead to reduced or inhibited growth in merozoites that managed to invade a new RBC. We found that there was no difference between the number of early ring stage parasites that progress through to late trophozoite stages (36 hours post-invasion) between GLCN treated and untreated cultures, indicating that PfCERLI1 knockdown merozoites that do invade do not have reduced viability during in-cycle growth. (Supplementary Fig. 2b). The similarity between growth and invasion inhibition, the normal development of merozoites and the build-up of free merozoites in the culture media strongly supports a direct role for PfCERLI1 in erythrocyte invasion.

### PfCERLI1 localises to the bulb of the rhoptry secretory organelle

Confocal microscopy of immune-labelled schizonts was used to assess the spatial positioning of HA-tagged PfCERLI1 relative to the micronemal marker Cysteine-rich protective antigen (CyRPA), inner membrane complex marker glideosome-associated protein 45 (GAP45), rhoptry neck marker rhoptry neck protein 4 (RON4) and rhoptry bulb marker rhoptry-associated protein 1 (RAP1) using confocal microscopy (Fig. 3a). Using thresholded Pearson’s correlation coefficient to calculate relative spatial proximity between compartments, semi-quantitative colocalisation analysis indicated that the signal of PfCERLI1 was associated most frequently with the rhoptry bulb marker RAP1 (Fig. 3b). Given the resolution limits of conventional confocal microscopy (~200 nm XY and ~500 nm Z)^31^ relative to the size of the double-bulbous rhoptries^19,32^, this approach was deemed unsuitable for the analysis of PfCERLI1 subcellular compartmentalisation within the rhoptries. To overcome this, we utilised Airyscan super-resolution microscopy, which revealed both PfCERLI1 and RAP1 displaying a double doughnut-shaped bulbous structure (Fig. 3c). This fluorescence pattern is analogous to the canonical structure of the rhoptry bulb and, combined with the close colocalisation between PfCERLI1 and RAP1 suggested that PfCERLI1 localises on or within the rhoptry bulb.

**Fig 3.**
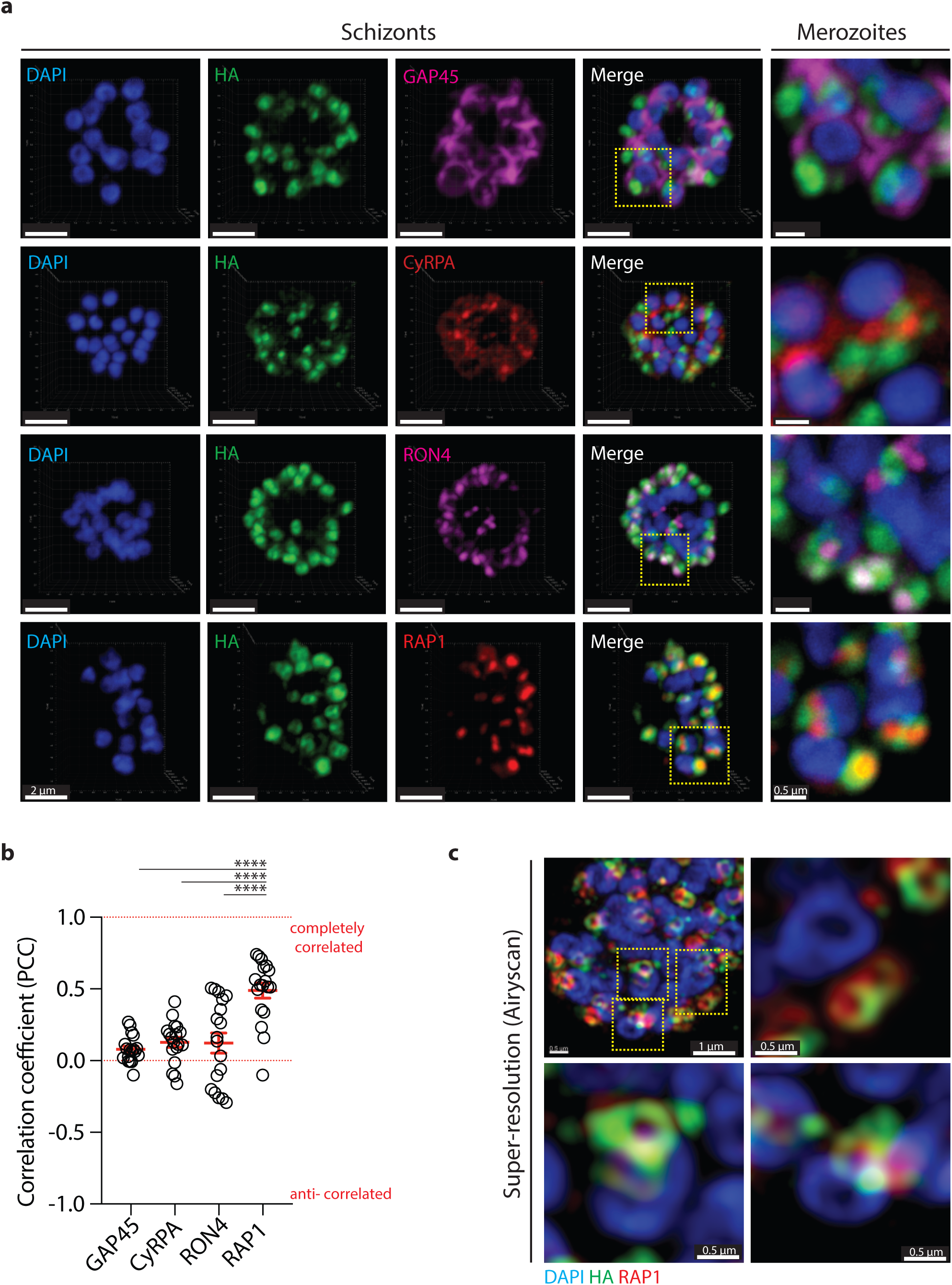
PfCERLI1 localises to the rhoptry bulb of merozoites. **(a)** Immunofluorescence microscopy showing 3D reconstructed projections of confocal images with anti-HA (PfCERLI1) in green, colocalising more strongly with the rhoptry bulb marker RAP1 than with the micronemal marker CyRPA, rhoptry neck marker RON4 or the inner-membrane complex marker GAP45. Scale bar = 1 μm. **(b)** Quantification of the colocalisation, by Pearson’s correlation coefficient when PfCERLI1 (anti-HA) staining is defined as the region of interest, with the following merozoite organelle markers: GAP45, CyRPA, RON4, and RAP1. (**** = p<0.0001 by Analysis of variance (ANOVA), n=3 treatments with 6 images from each biological replicate, error bars = SEM). **(c)** Maximum intensity projections of super-resolution immunofluorescence microscopy (Airyscan) of PfCERLI1^HAGlmS^ schizonts stained with anti-HA and anti-RAP1 antibodies.

In order to assess whether PfCERLI1 is on the inside or the outside of the rhoptry bulb, we biochemically characterised the location of this protein in relation to the membrane of the rhoptry organelle using a proteinase K protection assay. This assay determines the subcellular location of a protein as determined by its sensitivity to proteinase K after selective permeabilisation of non-organellar membranes with detergents. Since the rhoptry membrane is resistant to the detergent digitonin, proteins that lie inside the rhoptry are protected from proteinase K cleavage. Digitonin lysis of parasite membranes resulted in proteinase K cleavage of PfCERLI1^HAGlmS^ as detected by Western blot (Fig. 4a), indicating that PfCERLI1 lies outside the rhoptry and is exposed to the parasite cytosol. By contrast, RAP1, was not degraded significantly by proteinase K treatment, confirming that proteins inside the rhoptry lumen are refractory to digestion. Despite the level of colocalisation between PfCERLI1 and RAP1, this difference in proteinase K sensitivity suggested that PfCERLI1 localised to the cytosolic face of the rhoptry bulb.

**Fig 4.**
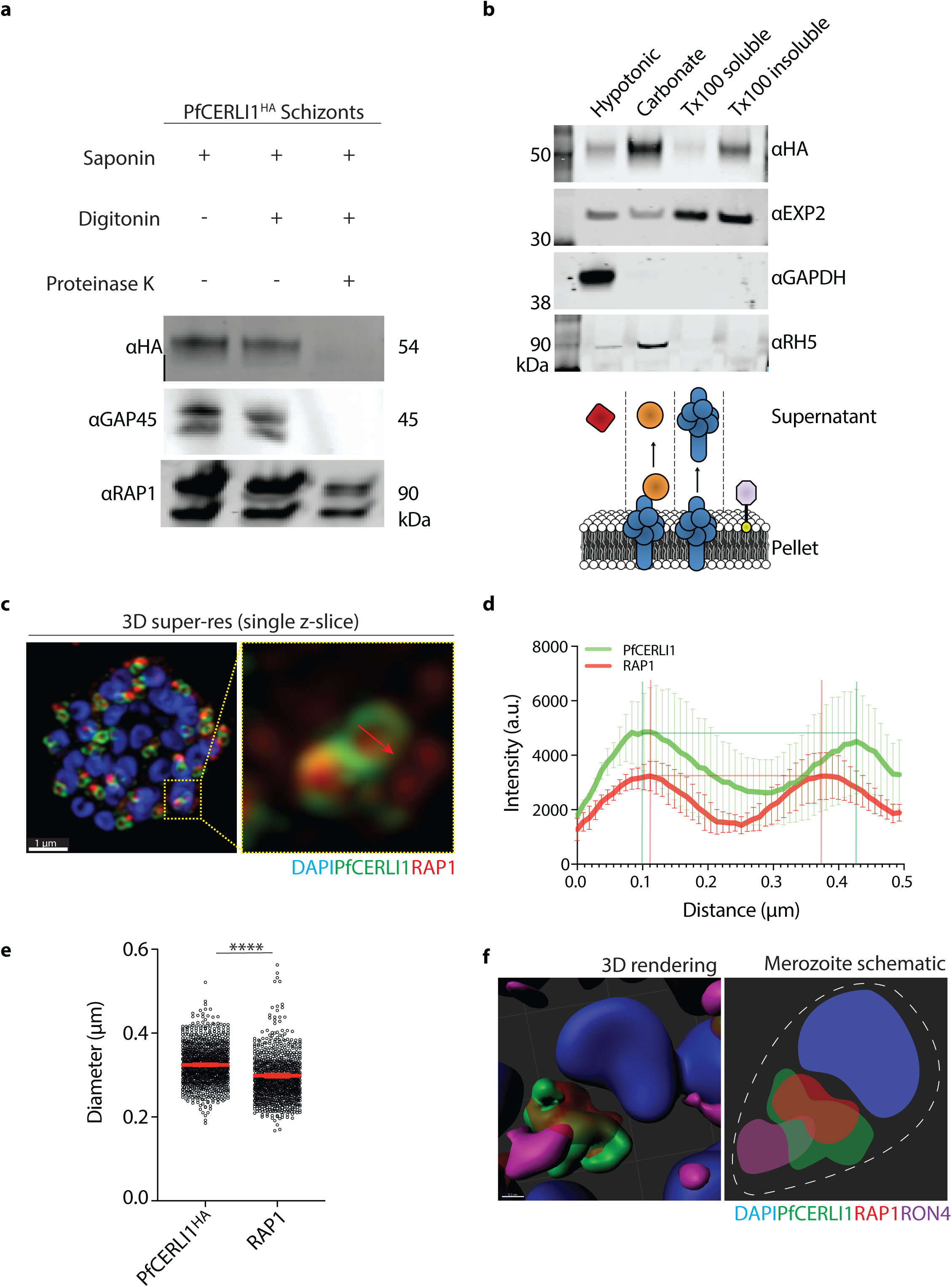
PfCERLI1 is peripherally-associated with the cytosolic face of the rhoptry membrane. **(a)** PfCERLI1^HA^ schizont cultures were used for a proteinase K protection assay. Parasites were treated with either saponin alone, saponin and digitonin or saponin, digitonin and proteinase K and probed with anti-HA antibodies (PfCERLI1). GAP45 (inner-membrane complex, exposed to the cytosol) and RAP1 (rhoptry lumen) serving as positive and negative controls, respectively, for proteinase K digestion. Selected images representative of three independent experiments. **(b)** To determine membrane association, PfCERLI1^HA^ schizont cultures were subjected to a solubility assay. Saponin lysed parasite cultures were hypotonically lysed before being treated with sodium carbonate and then Triton-X-100 (Tx100), with supernatants and the Tx100 insoluble fraction being reserved after each treatment. Resulting samples were probed with anti-HA antibodies (PfCERLI1). The presence of a strong band in the carbonate treatment indicates release of the majority of PfCERLI1 protein into the supernatant with this treatment. Membranes were also stained with anti-EXP2 (transmembrane domain containing), anti-GAPDH (cytosolic) and anti-RH5 (peripheral) solubility controls. Selected images representative of three independent experiments. **(c)** A single z-slice of the double-bulbous rhoptries displaying the scheme that was used to measure the fluorescence intensity peaks **(d)** and diameter **(e)** of anti-HA and anti-RAP1 staining across the rhoptry. **(f)** 3D rendered image of a free merozoite showing the nucleus (DAPI) localising at the basal surface, PfCERLI1 (HA) wrapping around RAP1 at the rhoptry bulb and RON4 localising in the rhoptry neck at the far apical tip.

Having identified that PfCERLI1 lies on the cytosolic face of the rhoptry bulb membrane, we assessed how HA-tagged PfCERLI1 was bound to the rhoptry membrane through sequential, differential solubilisation of lysed parasite membranes. Saponin pellets from PfCERLI1^HAGlmS^ schizont stage parasites were hypotonically lysed (to release non-membrane associated proteins), treated with sodium carbonate (Na_2_CO_3_; to release peripheral membrane proteins) and Triton X-100 (Tx100; to release integral membrane proteins). The supernatants containing solubilised proteins with the different treatments as well as the final Tx100 insoluble fraction (containing covalently lipid-linked proteins^33,34^) were used in Western blots with released PfCERLI1^HA^ protein detected using anti-HA antibodies (Fig. 4b). PfCERLI1 was detected primarily in the carbonate treatment, indicating that PfCERLI1 is likely to be peripherally associated with the cytosolic face of the rhoptry bulb membrane.

### PfCERLI1 is closely juxtaposed to the rhoptry bulb maker RAP1

Since biochemical analysis showed that PfCERLI1 associated with the cytosolic face of the rhoptry membrane whereas RAP1 appears confined to the rhoptry lumen, we used super-resolution microscopy to quantify the degree of physical proximity between PfCERLI1 and RAP1. To do this, we measured the radial positioning of all voxels that define the compartments of interest and plotted the signal intensities for both PfCERLI1 and RAP1 against the length of a vector drawn through the image (Fig. 4c). Using a line scan through a two-dimensional image of fluorescent objects, the distance between PfCERLI1 intensity peaks were significantly wider than that of RAP1 (Fig. 4d-e); further supporting the notion that RAP1 and PfCERLI1 are spatially segregated.

In order to assess whether PfCERLI1 is also associated with markers of the rhoptry neck that lie closer to the apical tip of the merozoite, we triple antibody-labelled compound 1 treated schizonts with anti-HA (PfCERLI1), anti-RAP1 (rhoptry bulb), and anti-RON4 (rhoptry neck) antibodies prior to super-resolution microscopy. Enhanced spatial resolution of fully-formed merozoites revealed that RON4 is located closest to the apical tip of the merozoite (Fig. 4f), as expected. Neither PfCERLI1 nor RAP1 exhibited extensive overlap with RON4, with the RAP1 signal enclosed entirely within that of PfCERLI1. These data support that PfCERLI1 lies on the outside of the rhoptry membrane closely juxtaposed to the rhoptry bulb marker RAP1 in the rhoptry lumen, confirming that PfCERLI1 is most closely associated with the outer membrane of rhoptry bulb in mature merozoites.

### PfCERLI1 contains domains that are involved with lipid-binding

In order to investigate how PfCERLI1 may be binding to the surface of the rhoptry bulb and provide insights on protein function based on solved protein structures, we used protein structural prediction and modelling software Phyre2^35^, I-TASSER^36–38^ and COACH^39,40^ to analyse PfCERLI1 structure. These analyses suggested that PfCERLI1 contains a C2 domain which are known lipid binding regions (amino acids ~50-170), a pleckstrin homology (PH) domain that typically binds calcium (amino acids ~250-360; Supplementary fig. 3a) and an N-terminal alpha helix that is currently annotated as a signal peptide. Outside of these two domains PfCERLI1 was predicted to be partially disordered. Like most canonical C2 domains^41^, the PfCERLI1 C2 domain was predicted to bind ionic calcium and phospholipids (Supplementary Fig. 3b). Of the models tested against, the PfCERLI1 C2 domain was most structurally similar to the C2 domain of human itchy homolog E3 ubiquitin protein ligase (ITCH) but the ligand(s) of this protein have not been elucidated^42^. The most structurally similar protein model the C2 domain of PfCERLI1 was that of C2-domain ABA-related protein 1 (CAR1) from *Arabidopsis thaliana*, which has been shown bind phospholipids in a manner that is dependent on a single calcium ion^43,44^. The PH domain of PfCERLI1 was most structurally similar to the PH domain of Protein kinase B/Akt, and like Akt, was predicted to bind to inositol 1,3,4,5-tetrakisphosphate (IP4) (Supplementary Fig 3c). IP4 forms the head-group of the phospholipid phosphatidylinositol (3,4,5)-trisphosphate (PIP_3_), which commonly coordinates protein membrane localisation^45,46^. Additionally, PfCERLI1 has previously been predicted to be palmitoylated^21^. Collectively, this suggests that PfCERLI1 contains potentially three different lipid-interacting moieties that may work cooperatively or independently to coordinate the binding of PfCERLI1 to the rhoptry bulb membrane.

PfCERLI1 residing on the cytosolic face of the rhoptry bulb membrane appeared to contrast with its predicted signal peptide, as golgi-mediated trafficking of signal peptide containing proteins should place them on the luminal side of golgi-derived organelles such as the rhoptries^32,47^. The putative signal sequence displayed on PlasmoDB^48^ is predicted using SignalP-3.0^49^, however, the latest version of this program (SignalP-5.0^50^) no longer predicts that PfCERLI1 contains an N-terminal signal sequence (Supplementary Fig. 4a). Conversely, RAP1, which we have demonstrated to be in the rhoptry lumen is predicted to have a signal peptide (Supplementary Fig. 4b). Considering the digestion of PfCERLI1 in the presence of proteinase K, we therefore hypothesise that this putative signal peptide does not lead to PfCERLI1 being trafficked through the secretory pathway via the golgi.

### PfCERLI1 knockdown alters rhoptry architecture but does not influence processing of invasion ligands

Having identified PfCERLI1’s confinement to the cytosolic face of the rhoptry bulb, we investigated whether PfCERLI1 may have a role in invasion ligand processing and distribution, or development of rhoptry architecture. Since a number of rhoptry antigens have been shown to be proteolytically cleaved inside the rhoptry lumen, we reasoned that if loss of PfCERLI1 function inhibited transport of rhoptry antigens to the lumen during biogenesis of the organelle, then we would be able to detect altered protein cleavage patterns in rhoptry antigens that undergo processing within this organelle. To assess this, GLCN treated and untreated PfCERLI1^HAGlmS^ schizont lysates were prepared for Western blot analysis. PfCERLI1 knockdown did not result in any consistent changes in processing of the rhoptry bulb marker RAP1, the rhoptry neck antigen RON4, or a control protein that is a component of the inner membrane complex (GAP45) that all undergo proteolytic cleavage inside the parasite prior to invasion commencing (Supplementary Fig. 1). Additionally, total protein levels of both RAP1 and RON4 remained consistent following PfCERLI1 knockdown suggesting there is no relationship between PfCERLI1 and total expression levels of these antigens in the parasite.

In order to investigate whether PfCERLI1 caused changes in rhoptry architecture, we visualised rhoptries using transmission electron microscopy (TEM) (Supplementary Fig. 5). Analysis of compound 1 matured schizonts failed to reveal any obvious ultrastructural differences between the rhoptries of either GLCN treated or untreated PfCERLI1^HAGlmS^ parasites.

While the qualitative EM assessment indicated that PfCERLI1 knockdown was not causing clearly discernible changes in rhoptry architecture, we investigated whether there was altered rhoptry antigen distribution with PfCERLI1 knockdown using semi-quantitative super-resolution microscopy. GLCN treated and untreated PfCERLI1^HAGlmS^ (Supplementary Fig. 6a) parasites were imaged in 3-dimensions to estimate shape (sphericity, oblate (flattened sphere), prolate (elongated sphere)), staining intensity, volume and area for the rhoptry bulb marker RAP1 (Supplementary Fig. 6b). A highly significant decrease in the intensity (50% reduction) and increase in the size of RAP1 staining with PfCERLI1 knockdown was observed across five biological replicates. These changes were associated with a significant elongation of the RAP1 signal, but no corresponding change in the RAP1 diameter, indicating that the change in RAP1 staining intensity and size is largely due to elongation of the protein’s distribution in the rhoptry (Fig. 5e). Collectively, these changes in intensity and size suggest that knockdown of PfCERLI1 alters the distribution of RAP1 within the rhoptries, and/or rhoptry shape more grossly.

**Fig 5.**
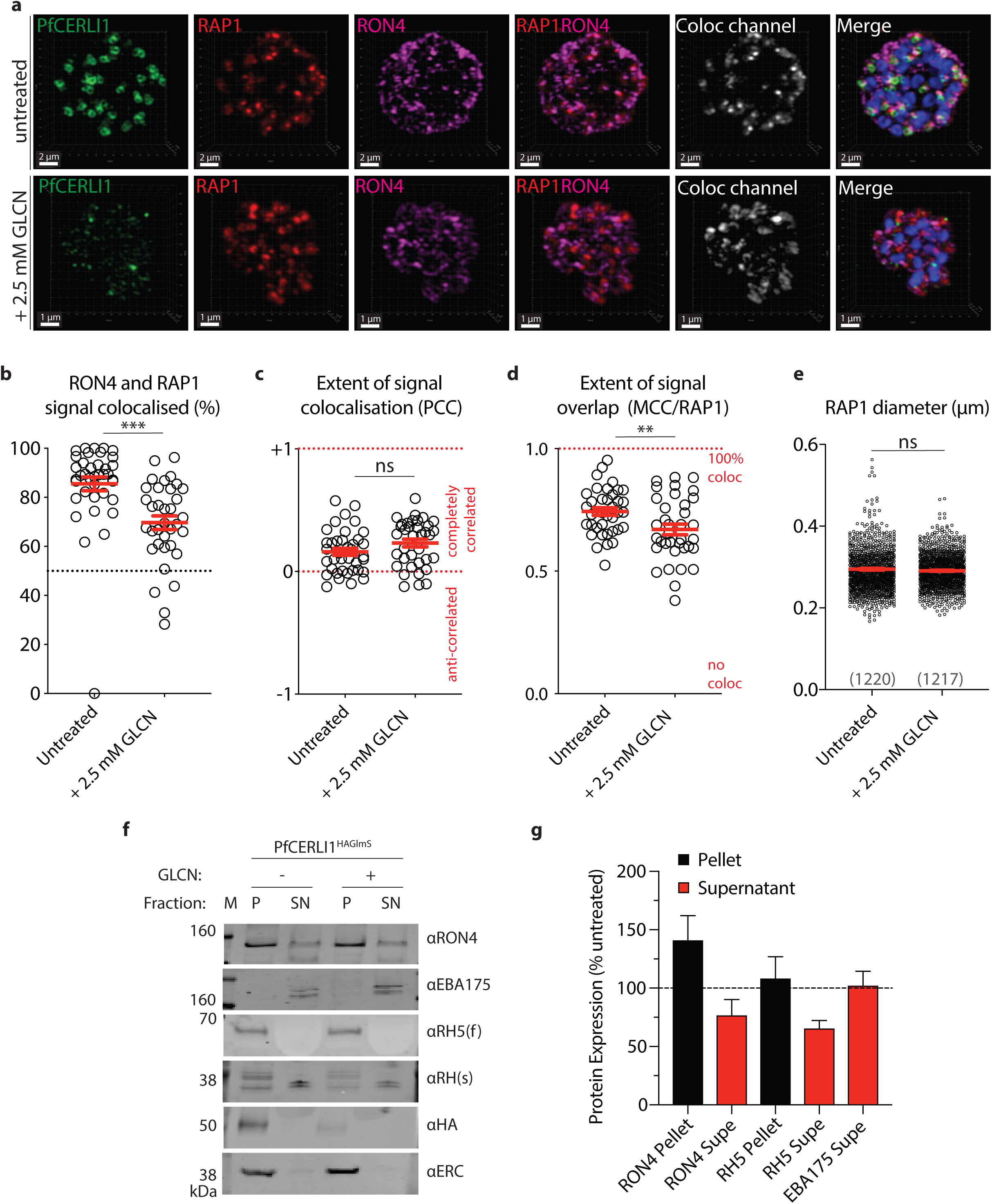
PfCERLI1^HAGlmS^ knockdown alters rhoptry antigen distribution and inhibits rhoptry secretion. **(a)** 3D reconstructions super-resolution (Airyscan) micrographs of PfCERLI1^HAGlmS^ schizont cultures that were either left untreated or treated with glucosamine (+2.5 mM GLCN) before being stained with DAPI (nucleus), anti-HA (PfCERLI1), anti-RAP1 (rhoptry bulb) and anti-RON4 (rhoptry neck) antibodies. According to the image analysis pipeline detailed in Supplementary Figure 7, the **(b)** % of RON4 that colocalised with RAP1, **(c)** Pearson’s correlation coefficient (PCC) of the amount of RON4 colocalising with the RAP1 signal, and **(d)** thresholded Mander’s correlation coefficient (MCC) of RON4 overlap with RAP1 signal was compared between PfCERLI1^HAGlmS^ knockdown and control parasites. Each data point represents a single schizont image (n=5 biological replicates, error bars = SEM). **(e)** Rhoptry bulb (RAP1) diameter was measured between untreated and PfCERLI1^HAGlmS^ knockdown (+2.5 mM GLCN) parasites. Each data point represents a single rhoptry (n=5 biological replicates, total number of rhoptries listed below each column). (ns = p>0.05, ** = p<0.01, *** = p<0.001 by one-way unpaired t-test). **(f)** PfCERLI1^HAGlmS^ parasites were either treated with 2.5 mM GLCN (+) or left untreated (-) and had surface receptors cleaved by enzyme treatment to prevent invasion but allow release of merozoite antigens. The parasite lysate (P) and culture supernatant (SN) were stained with anti-RON4 (rhoptry neck), anti-EBA175 (micronemes), anti-RH5 (rhoptry neck, f = full length, s = secreted), anti-HA (PfCERLI1) and anti-ERC (loading control) antibodies and analysed by western blot. Selected blots representative of five independent experiments (M = size markers). **(g)** Western blots of rhoptry antigens, and secreted EBA175, were normalised to the loading control (ERC) and quantified, with data represented as + 2.5 mM GLCN signal expressed as a % of the signal for the untreated control (error bars = SEM).

To further investigate the spatial positioning of rhoptry antigens, we quantified spatial proximity between RAP1 and RON4 markers in super-resolved GLCN treated and untreated PfCERLI1^HAGlmS^ schizonts (Fig. 5a-e). We first assessed whether distribution of RON4 changes with PfCERLI1 knockdown as described for RAP1. There was a trend towards greater RON4 intensity (12% increase), probably as a result of a less elongated signal in the rhoptry neck (Supplementary Fig. 6c,d). We next investigated whether changes in RON4 and RAP1 staining altered their spatial localisation relative to each other. Using Mander’s correlation coefficient to calculate association frequency between markers, semi-quantitative colocalisation analysis indicated that the proportion of RAP1 signal that overlapped with RON4 after knockdown of PfCERLI1 was significantly reduced (Fig. 5a-d). These data indicate that PfCERLI1 knockdown causes changes in the distribution of antigens within the rhoptry, most notably for RAP1, and increases the spatial segregation between rhoptry neck (RON4) and bulb (RAP1) markers.

### PfCERLI1 knockdown interferes with secretion of rhoptry antigens

After identifying that PfCERLI1 knockdown caused a significant change in the distribution of the rhoptry bulb protein RAP1, we next investigated whether knockdown of PfCERLI1 changed apical organelle secretion. Ring-stage cultures were either treated with 2.5 mM GLCN or left untreated before subsequent enzyme removal of erythrocyte surface receptors at trophozoite stages to prevent reinvasion. Merozoites were allowed to rupture, the supernatant collected, and lysates were made from saponin lysed parasite material before analysis by Western blot (Fig. 5f). Blots were probed with anti-RON4 and anti-RH5 antibodies to assess rhoptry secretion, anti-EBA175 to assess microneme secretion, anti-HA to assess PfCERLI1 knockdown and anti-ERC as a loading control. Knockdown of PfCERLI1 did not significantly alter the level of EBA175 (microneme) secretion into the supernatant relative to untreated controls, confirming that release of microneme contents had been triggered for both GLCN and untreated parasites (Fig. 5g). In contrast, quantitation of Western blots revealed a decrease in secreted RON4 and RH5 in the supernatant for PfCERLI1 knockdown parasites relative to untreated controls (Fig. 5g). Additionally, both RON4 and RH5 were observed to be more concentrated in the GLCN treated PfCERLI1^HAGlmS^ parasite pellet relative to untreated control parasite material. These data suggest that PfCERLI1 knockdown prevents secretion of essential invasion antigens from the apical tip of the rhoptry after initiation of the invasion process, providing a probable mechanism by which PfCERLI1 knockdown inhibits invasion.

## Discussion

Previous studies have identified PfCERLI1 (Pf3D7_0210600) as being a potential vaccine target on the basis of its predicted signal peptide and its stage of expression that peaks at late schizont, a transcription profile typical of a protein involved in merozoite invasion^23^. Additionally, PfCERLI1 was identified as a member of the ‘invadome’, an interaction network of putative invasion-associated proteins, and was localised to the merozoite apical tip in this study^22^. A random mutagenesis study that failed to recover viable parasites with disruption of PfCERLI1 indicates that this protein may have an essential role in parasite survival^25^. Characterisation of naturally acquired antibody levels towards PfCERLI1 using ELISA found only low-level antibody binding^23^. Indeed, there is no direct functional evidence showing localisation of PfCERLI1 on the outside of the merozoite where it could be targetable by immune responses and developed as a vaccine candidate. Here we show PfCERLI1 has an essential role in merozoite invasion, but also demonstrate that this protein lies on the cytosolic face of the rhoptry bulb membrane, a localisation that would prevent exposure to antibodies during the natural course of invasion.

The vast majority of previously characterised rhoptry proteins, including all members of both the high (RhopH) and low (RAP) molecular weight rhoptry complexes, rhoptry neck (RON) proteins and reticulocyte-binding family homologs (RH), possess a predicted signal peptide, allowing trafficking to the Golgi-derived nascent rhoptries^15,17,48,51–56^. There are notable exceptions to this, however, including Rhoptry protein (RHOP) 148 and armadillo-repeats only (ARO), neither of which contain a predicted signal peptide^57,58^. While the mechanism for rhoptry localisation of Rhop148 has not been explored, ARO localises to the cytosolic face of the rhoptry membrane through dual acylation of its N-terminus^58^. Another noteworthy protein is the recently described merozoite organising protein (MOP), which localised closely to, but not within, the rhoptry neck by both confocal and immuno-electron microscopy^59^. Additionally, MOP was shown to be cytosolically exposed by proteinase K protection assay. By contrast to ARO and RHOP148, MOP has no myristylation or palmitoylation sites and its localisation is hypothesised to be coordinated by protein-protein interactions^59^.

Here we show that PfCERLI1 localises to the rhoptries and is confined to the cytosolic face on the rhoptry bulb membrane. Semi-quantitative super-resolution fluorescence microscopy showed a difference between the mean diameter of PfCERLI1 and the rhoptry bulb marker RAP1 around the bulb structure, indicating they occupy distinct compartments on/within the rhoptry. Interestingly, we also observed that RAP1 consistently stained in the shape of a cylinder with an unstained core, suggesting that distribution of this protein in the lumen of the rhoptry bulb is not uniform and there may be internal structures that confine this protein around the internal diameter of the bulb. PfCERLI1 was sensitive to proteinase K after selective membrane solubilisation and RAP1 which is protected inside the rhoptry lumen was insensitive, confirming that PfCERLI1 compartmentalises to the cytosolic face of the rhoptry bulb membrane. The spatial separation of PfCERLI1 and RAP by microscopy and biochemical analysis may support the hypothesis that an internal or external scaffold is present that supports rhoptry structure, but resolution limits and the size of the antibodies conventionally used for immunofluorescence microscopy prevents more precise localisation. PfCERLI1 has been shown to be palmitoylated^21^, but was predominantly present in the carbonate fraction of the membrane solubility assay (Fig. 4b), suggesting that PfCERLI1 predominantly associates with the cytosolic face of the rhoptry membrane through protein-protein or protein-lipid electrostatic interactions in schizonts. PfCERLI1 was also detected in the Tx100 insoluble fraction. While the contents of this fraction are poorly understood in *Plasmodium*, they have been previously identified as containing covalently lipid-linked (palmitoylated and myristoylated) members of the IMC^60–62^. C2 domains, as present in PfCERLI1, are predicted to bind calcium and known to form ionic bridges with lipid phosphates in membrane bilayers that are not easily disruptable^63^, providing a second potential membrane binding site for PfCERLI1. The presence of two distinct moieties that are predicted to interact with (C2 domain and palmitoylated residues) may explain why PfCERLI1 partially solubilises in carbonate buffer while the remainder stays in the Tx100 insoluble fraction.

Our attempts to knock-out *Pfcerli1* proved unsuccessful, confirming a previous random mutagenesis study that identified this gene as essential to asexual parasite growth^25^. However, we successfully integrated a *GlmS*-ribozyme sequence at the 3’ end of the *Pfcerli1* gDNA sequence and were able to knockdown protein expression by >80% with the addition of GLCN for periods as short as 24 hours during merozoite development. Using this system, knockdown of PfCERLI1 protein expression led to >45% inhibition of parasite growth. While we were unable to achieve complete growth inhibition, it is possible that loss of PfCERLI1 function is compensated for by other proteins with similar function. Indeed, bioinformatic searches identified a second protein, Pf3D7_0405200 (now called PfCERLI2), with similar structure and shared predicted epitopes that has previously been implicated in merozoite invasion^64^, providing a possible functional homologue that could compensate for loss of PfCERLI1.

Knock-out and knockdown studies implicate PfCERLI1 as having an important role in asexual stage parasite growth and we further confirmed that loss of this protein led to a defect in invasion. Initial investigation of the PfCERLI1 knockdown parasites showed that the number and morphology of merozoites was unaffected by loss of PfCERLI1 and parasites that did invade with PfCERLI1 knockdown developed normally. We determined that PfCERLI1 knockdown led directly to a significant reduction in newly invaded ring stage parasites and a proportional build up in free merozoites in the culture, confirming that knockdown of PfCERLI1 inhibits merozoite invasion directly prior to the formation of the irreversible tight junction, which would ordinarily bind the merozoites to the erythrocyte surface. Given that inhibition due to PfCERLI1 knockdown was almost identical between growth and invasion assays, invasion inhibition is likely the exclusive cause of knockdown-induced growth inhibition and therefore implicates an essential role for PfCERLI1 in merozoite invasion.

When PfCERLI1 was knocked down, we saw evidence of aberrant rhoptry antigen distribution, but not antigen processing, suggesting that this protein has either a structural or mechanical role required for release of rhoptry contents, rather than a significant role in transporting in unprocessed rhoptry antigens to the rhoptries. Semi-quantitative super-resolution fluorescence microscopy revealed a significant decrease in RAP1 staining intensity with PfCERLI1 knockdown which corresponded to increases in RAP1 volume and area, indicating knockdown of PfCERLI1 changes RAP1 distribution. While changes in area and volume for RAP1 were observed, the diameter of RAP1 rhoptry bulb staining and TEM analysis indicated that there were no large morphological changes in the rhoptry bulb structure itself. Thus, changes in the ultrastructure of the rhoptry with PfCERLI1 knockdown appear to be minimal and the strongest indication of a modified rhoptry shape achieved to date is increase prolate staining for RAP1, which may indicate a lengthening of the rhoptries that was not noticeable in the single sections in the TEM images.

To assess whether the segregation of rhoptry antigens was affected by loss of PfCERLI1, we compared the colocalisation of RAP1 (bulb) and RON4 (neck) by super-resolution microscopy. Knockdown of PfCERLI1 led to a significant decrease in the proportion of RAP1 overlapping with RON4. With PfCERLI1 knockdown leading to invasion inhibition and changes to rhoptry antigen distribution, we assessed whether knockdown altered secretion of rhoptry antigens during the invasion process. Using apical organelle secretion assays, we identified that knockdown of PfCERLI1 resulted in a decrease in the secretion of the rhoptry antigens RH5 and RON4 but no major change in secretion of the micronemal antigen EBA175. In addition, the reduction of RON4 in the supernatant corresponded to an increase in RON4 in the parasite pellet, suggesting that release of RON4 was inhibited after initiation of the invasion process. As antigens secreted from the rhoptry neck are essential for merozoite invasion^16,65^, disruption of rhoptry neck antigen secretion withPfCERLI1 knockdown points to a role for this protein in successful rhoptry antigen secretion during the invasion process.

Here, we describe a key role for PfCERLI1 in controlling the distribution and secretion of rhoptry antigens during the essential step of merozoite invasion into the host erythrocyte. Identification of PfCERLI1’s direct association with release of rhoptry antigens is the first step in understanding the complex molecular events that control rhoptry secretion during invasion. By understanding how rhoptry secretion is controlled and the key proteins involved, we can identify targets for drug development that will stop merozoites invading and prevent replication of disease-causing parasites. Additionally, this study makes extensive use of semi-automated quantitative immunofluorescence microscopy and highlights how this powerful tool can be used to study the process of invasion.

## Methods

### Bioinformatic analyses

PfCERLI1 (Pf3D7_0210600) and orthologous sequences in *P. reichenow*i (PRCDC_0209500), *P. gaboni* (PGAB01_0208200), *P. malariae* (PmUG01_04021700), *P. ovale* (PocGH01_04019500), *P. knowlesi* (PKNH_0410600), *P. vivax* (PVX_003980), *P. berghei* (PBANKA_0307500), *P. yoelii* (PY17X_0308100) and *P. chabaudi* (PCHAS_0309700) were obtained by searching within the PlasmoDB.org database^48^. Sequence similarities were determined using Geneious 9.1.3 (Biomatters) and performing multiple pairwise alignments using the global alignment with free end gaps alignment algorithm with the Blosum62 cost matrix.

### Continuous culture of asexual stage *P. falciparum*

*P. falciparum* (3D7, 3D7 Δ*Pfcerli1*^HAGlmS^ & 3D7 Δ*Pfcerli1*^HAGlmS/GFP^) parasites were cultured in human O^+^ erythrocytes (Australian Red Cross blood service) as previously described^66^. Parasites were grown in RPMI-HEPES culture medium at pH 7.4 (Gibco) that was supplemented with 50 μM hypoxanthine, 25 mM NaHCO_3_, 20 μM gentamicin and either 0.5% Albumax II (Gibco) or 0.25% w/v Albumax II, 5% v/v human serum (Australian Red Cross blood service). Parasite cultures were maintained in an atmosphere of 1% O_2_, 4% CO_2_ and 95% N_2_ at 37 °C.

### Plasmid construction and transfection

The PfCERLI1^HAGlmS^ riboswitch transfection vector was prepared from the PTEX150^HAglmS^ vector^26,67^. The final 767 base pairs of the 3’ end of the *Pfcerli1* genomic sequence (excluding the stop codon) was PCR amplified using the primers *Pfcerli1* 5’ F RBW (GGT**AGATCT**CATATCAAATTTGGTTCTTGAAG) and *Pfcerli1* 3’ R RBW (GGT**CTGCAGC**ATCACTATAGTTGTACATATTTTTGC). The resulting PCR fragment was cloned into the PTEX150-^HAglmS^ vector using the restriction enzyme cloning sites Bgl II and Pst I (restriction enzyme sites in bold). To generate the cytosolic GFP expressing 3D7 Δ*Pfcerli1*^HAGlmS/GFP^ line, the pHGBrHrBl-1/2 GFP plasmid was used, without modification, as previously described^29^. For PfCERLI1 disruption, a source vector (Pf3D7_1463000 SLI-TGD) was digested with Not I and Mlu I with a 5’ *Pfcerli1* flank being amplified using the primers (restriction enzyme sites in bold) *Pfcerli1* SLI-TGD F (GGT**GCGGCCGC**GATACTCACAACATATTATATCTTGG) and *Pfcerli1* SLI-TGD R (GGT**ACGCGT**CATACCTCTATGTGTACTTTGTTCTG)^24^.

*P. falciparum* 3D7 parasites were transfected using erythrocyte loading^68^. Briefly, uninfected erythrocytes were centrifuged at 1500 rcf for 1 minute, before removal of the supernatant and washing in cytomix (0.895% KCl, 0.0017% CaCl_2_, 0.076% EGTA, 0.102% MgCl_2_, 0.0871% K_2_HPO_4_, 0.068% KH_2_PO_4_, 0.708% HEPES). Erythrocytes were then resuspended in cytomix containing 200 μg of ethanol precipitated plasmid DNA, incubated in a 0.2 cm cuvette (Bio-Rad) on ice for 30 minutes and then electroporated (Bio-Rad) at 0.31 kV with a capacitance of 960 μF. Cells were washed 2X with culture media and transferred to a 10 mL dish containing magnet purified schizont stage parasites (3-5% parasitaemia). Integration of the HA-*GlmS* plasmid was selected for using 3 cycles of 5 nM WR99210 (Jacobus Pharmaceuticals) drug treatment. For generation of the Δ*Pfcerli1*^HAGlmS/GFP^ line, PfCERLI1^HAGlmS^ parasites were transfected with the an episomal cytosolic GFP expressing plasmid pHGBrHrBl-1/2 and maintenance of this plasmid was selected for using 5 μg/mL blasticidin-S-deaminase HCl (Merck Millipore). Maintenance of the SLI-TGD plasmid was selected for using 20 μM WR99210, with successful integrants then selected for using 400 μg/mL G418 sulphate (geneticin, Thermo Fisher).

### Assessment of *in vitro* blood stage growth and invasion

To determine the number of merozoites produced by each fully formed schizont after knockdown of PfCERLI1 protein expression, thin blood smears were made of D-(+)-Glucosamine hydrochloride (Sigma-Aldrich) (GLCN) treated and untreated ate schizont stage cultures matured in the presence of E64 (Sigma-Aldrich) to prevent schizont rupture. Smears were methanol fixed and stained with Giemsa (Merck Millipore) before blinded assessment of the number of merozoites by light microscopy (20 individual schizonts). All parasites assessed were fully matured, with individual segmented merozoites visible.

To assess the impact of PfCERLI1 knockdown, 3D7 Δ*Pfcerli1*^HAGlmS^ (growth) and 3D7 Δ*Pfcerli1*^HAGlmS/GFP^ (invasion) parasites were synchronised to ring stages using sorbitol lysis and assays were set up in 96-well U-bottom plates at 1% parasitaemia and 1% haematocrit in a volume of 45 μL as described previously^69^. 5 μL of 10x concentration GLCN or complete media was added to make a final volume of 50 μL. Assays were stained with 10 μg/mL ethidium bromide (Bio-Rad) in PBS before assessment of parasitaemia using flow-cytometry (BD Biosciences LSR II, 488 nm laser with FITC and PE filters). To assess merozoite development and invasion of 3D7 Δ*Pfcerli1*^HAGlmS/GFP^ parasites, GLCN treated and untreated cultures were grown for 36 hours, until newly invaded rings were present (0-6 hours post-invasion). Flow cytometry data was analysed using FlowJo (Tree Star).

### Saponin lysis and Western blot

For protein samples, ~10 mL of high parasitaemia culture was lysed with 0.15% w/v saponin for 10 minutes on ice, parasite material was pelleted by centrifugation before washing once in 0.075% w/v saponin and three times in PBS. In the presence of protease inhibitors (CØmplete, Roche). Parasite lysates were resuspended in reducing sample buffer (0.125 M Tris-HCl pH 7, 4% v/v SDS, 20% v/v glycerol, 10% v/v β-mercaptoethanol (Sigma-Aldrich), 0.002% w/v bromophenol blue (Sigma-Aldrich)) and separated by size using SDS-PAGE 4-12% Bis-Tris Gels (Bolt, Invitrogen) at 130 V for 60 min. Proteins were then transferred to a nitrocellulose membrane (iBlot, Invitrogen) at 20 V for 7 minutes, before blocking the membrane for one hour at room temperature in Odyssey Blocking Buffer (TBS) (LI-COR Biosciences). Primary (mouse 12CA5 anti-HA (1:4000, Roche), rabbit anti-EXP2 (1:5000,^70^), rabbit anti-GAP45 (1:10000,^71^), mouse anti-RAP1 (1:5000,^72^), rabbit anti-RON4 (1:5000), rabbit anti-EBA175 (1:5000^73^), mouse anti-RH5 (1:5000 **REF**), rabbit anti-ERC (1:10000^74^) and secondary (IRDye ® 800CW goat anti-mouse (1:4000, LI-COR Biosciences), IRDye ® 680RD goat anti-rabbit (1:4000, LI-COR Biosciences)) antibodies were incubated with membranes for one hour at room temperature (prepared in Odyssey Blocking Buffer (TBS)), before washing in 0.05% v/v PBS Tween followed by a PBS wash after the secondary antibody incubation. Western blots were visualised using an Odyssey Infrared imaging system (LI-COR Biosciences). Western blot quantification was performed using Image Studio Lite 5.2.5 (LI-COR Biosciences).

### Proteinase K protection assay

Proteinase protection assays were modified from a previously published study^58^. Three 5 mL aliquots of high-parasitaemia E64 treated schizonts were centrifuged at 440 rcf for 5 minutes before removal of supernatant and washing in 1xPBS and lysing of uninfected RBCs using saponin. The following treatments were undertaken: One tube with SOTE buffer (0.6 M sorbitol, 20 mM Tris HCl pH 7.5, 2 mM EDTA) alone. A second tube with SOTE plus 0.02% w/v digitonin (Sigma-Aldrich) incubated for 10 minutes at 4 °C prior to washing in SOTE buffer. A third tube with digitonin treatment followed by digestion with 0.1 μg/μL Proteinase K (Sigma-Aldrich) in SOTE for 30 minutes at 4 °C. Proteinase K was inactivated by adding 50 μL of 100% v/v trichloroacetic acid followed by a PBS wash. All tubes were then resuspended in 500 μL acetone before cells were pelleted, the supernatant removed, and the pellet used for Western blot analysis of proteinase K sensitivity.

### Protein solubility assay

Protein solubility assays were performed as previously described^58^. Briefly, saponin-lysed pellets from 10 mL of high-parasitaemia E64 treated schizonts were resuspended in 100 μL dH_2_O, snap-frozen at −80 °C four times, passed through a 26-gauge needle 5 times to disrupt the parasite membrane and then centrifuged at 15,000 rcf for 10 minutes with the water-soluble supernatant reserved. The pellet was washed twice in dH2O and once in 1xPBS before resuspension in 0.1M sodium carbonate (Na_2_CO_3_) for 30 minutes at 4 °C with the supernatant reserved. Following carbonate treatment, samples were resuspended in 0.1% v/v Triton-X-100 for 30 minutes at 4 °C with the supernatant reserved and the resulting pellet washed and resuspended in 1 x PBS. Samples were analysed by Western blotting.

### Sample preparation for microscopy

Untreated and GLCN treated cultures (approximately 10 mL of 3% parasiteamia) of Compound 1 arrested PfCERLI1^HA*GlmS*^ late-stage schizonts were concentrated by centrifugation at 1700 rpm for 3 minutes, washed in PBS and fixed with 4% v/v paraformaledehyde (PFA, Sigma-Aldrich), 0.0075% v/v glutaraldehyde (pH 7.5, Electron Microscopy Sciences) solution for 30 minutes at room temperature with gentle shaking. Fixed parasite suspensions were adjusted to 1% haematocrit prior to adhering onto 0.01% poly-L-lysine (Sigma-Aldrich) coated #1.5H high-precision coverslips (Carl Zeiss, Oberkochen, Germany) for 1 hour at room temperature before permeabilistion with 0.1 % v/v Triton-X-100 for 10 minutes. Coverslips were incubated with fresh blocking solution (3% w/v BSA-PBS, 0.05% v/v Tween-20) for at least 1 hour. Primary antibodies (Chicken anti-HA (Abcam), mouse anti-RAP1^72^, mouse anti-CyRPA 8A7^75^, rabbit anti-RON4^76^, rabbit anti-GAP45^71^) were diluted 1:500 in antibody diluent (1% w/v BSA-PBS, 0.05% v/v Tween-20) were added to coverslips for 1 hour at room temperature of overnight at 4°C. Cells were washed three times with PBS-Tween-20 (0.1 % v/v) and incubated with goat anti-chicken/mouse/rabbit Alexa Fluor coupled secondary antibodies (488 nm, 594 nm, 647 nm) (Life Technologies) diluted 1:500 for 1 hour at room temperature. After secondary antibody incubations, coverslips were washed three times with PBS-Tween-20 (0.1% v/v) then post-fixed with 4% w/v PFA for 5 minutes. PFA was washed off and coverslips dehydrated in ethanol at 70% v/v (3 minutes), 95% v/v (2 minutes) and 100% v/v (2 minutes) before being allowed to air dry and mounted on slides with ProLong® Gold antifade solution (refractive index 1.4) containing 4’, 6-diamidino-2pheynlindole, dihydrochloride (DAPI) (ThermoFisher Scientific). Once the moutant had cured for 24 hours, cells were visualised by either conventional confocal (Olympus FV3000) or Zeiss LSM 800 Airyscan super-resolution microscopy (Carl Zeiss, Obekochen, Germany).

### Confocal microscopy

Conventional confocal microscopy was performed on Olympus FV3000 fluorescent microscope (Olympus) equipped with a 100X MPLAPON oil objective (NA 1.4) using the 405 nm, 488 nm, 561 nm and 633 nm lasers. Z-stacks were acquired with a step size of 0.43 μm using a sequential scan (scan zoom = 10, without line averaging).

### Super-resolution microscopy

Sub-diffraction microscopy was performed on a Zeiss LSM800 AxioObserver Z1 microscopy (Carl Zeiss, Obekochen, Germany) fitted with an Airyscan detector and a Plan-Apochromat 63X (NA 1.4) M27 oil objective. The TetraSpeck™ Fluorescent Microspheres (Life Technologies) were mounted on #1.5H coverslips (Carl Zeiss, Oberkochen, Germany) with a density of ~2.3 × 10^10^ particles/ml and used as the 200 nm bead calibration sample to correct for chromatic and spherical aberrations. The super-resolution reconstructions of multi-labelled PfCERLI1^HA*GlmS*^ schizonts were acquired, sequentially in four channels, as follows: channel 1 = 633 nm laser, channel 2 = 561 nm laser, channel 3 = 488 nm laser, channel 4 = 405 nm laser, with exposure times optimised from positive and negative control samples (HA blocking peptide, or omitted primary antibody) to avoid saturation. Three-dimensional (3D) Z-stacks were acquired at a pixel resolution of 0.04 μm in XY and 0.16 μm intervals in Z using piezo drive prior to being Airyscan processed in 3D using batch mode in ZEN Black (Zeiss).

### Diameter measurements of super-resolved rhoptries

For further spatial exploration of the colocalised signals, 3D volumes captured with super-resolution microscopy after Airyscan processing were imported into ZEN Blue (Zeiss) for object-based colocalisation analysis. In this approach, volumetric colocalisation relies on manual identification of structures of interest and a subsequent measurement of their fluorescence intensity curves. Each rhoptry bulb immunolabelled with anti-RAP1 and anti-HA antibodies was manually quantified by a researcher blinded to experimental conditions, and the organelle diameter measured by drawing a vector through the centre of these structures and plotting the fluorescence intensities for the green and red channel against the length of the vector. Fluorescence intensity profiles of overlapping subcellular structures were then analysed in successive single sections from an image stack representing the two RAP1 and PfCERLI1^HAGlmS^ labelled structures. Fluorescence curves between the two antipodal points on the surface of the ring-like organelle were exported as .csv Excel files and plotted against the distance in nanometres. Only rhoptries with homogeneously stained ring-like structures containing no gaps in RAP1 or PfCERLI1^HAGlmS^ labelling and >100 nm in size were used for diameter measurements. Schizonts densely packed with merozoites with overlapping rhoptries were excluded from scoring. More than 800 individual organelles were scored and data from biological replicates plotted in Prism for comparison of diameters between untreated and glucosamine treated parasites.

### Rhoptry and microneme secretion assays

Apical organelle secretion assays were modified from a previous protocol^59^. Briefly, synchronous ring-stage cultures were obtained using sorbitol and either treated with 2.5 mM GLCN or left untreated and incubated in standard culture conditions for 24 hours until early trophozoite stage. Cultures were then enzyme treated with neuraminidase (0.067 U/mL), chymotrypsin (1 mg/mL) and trypsin (1 mg/mL). Parasites were then allowed to rupture over the following 24 hours. Following rupture, cultures were centrifuged at 13,000 rcf for 10 minutes in a benchtop centrifuge with some culture supernatant reserved, stored on ice, and the rest removed. The remaining pellet was then also placed on ice and saponin lysed to form a lysate containing the parasites that did not rupture and free merozoites, with this pellet and the culture supernatant analysed by Western blot.

### Statistical analysis

Graphs and statistical analyses were performed using GraphPad PRISM 7 (GraphPad Software Inc.). In all figures where p-values were calculated, the corresponding statistical test is listed in the figure legend along with the number of experiments results are pooled from.

## Acknowledgements

We thank Prof. Alan Cowman for provision of CyRPA, RH5, RON4, EBA175 and GAP45 antibodies, Dr. Paul Gilson for EXP2 antibodies and Prof. Leann Tilley for ERC and GAPDH antibodies. We also thank Dr. Paul Gilson for the PTEX150^HAGlmS^ transfection vector. Confocal microscopy was performed at Adelaide Microscopy, University of Adelaide, and super-resolution microscopy was performed at the Centre for Cancer Biology Cytometry Facility, The University of South Australia. We especially thank Dr. Jane Sibbons from Adelaide Microscopy for assistance with microscopy. Electron microscopy was performed at the Bio21 Institute Advanced Microscopy Facility, The University of Melbourne (www.microscopy.unimelb.edu.au). For provision of the SLI-TGD vector, we thank Dr. Tobias Spielmann. We thank Dr. Brad Sleebs for Compound 1. We thank Arne Alder and Sarah Lemcke for help with SLI-TGD transfection and parasite culture.

This work was supported by funding from the NHMRC (Project Grant APP1143974, DW), University of Adelaide Beacon Fellowship (DW), DAAD/Universities Australia joint research co-operation scheme (TG, DW, BL), Australian Government Research Training Program Scholarship (BL), South Australian Commonwealth Scholarship (BL).

## Author contributions

Study design and planning: D.W.W., B.L., S.F., M.W.A.D., and T.G. Performed experiments and generated reagents: B.L., S.F., D.W.W., G.K.H. and B.Liu. Data analysis: S.F, B.L. and D.W.W. Manuscript writing: B.L, S.F and D.W.W. Manuscript was drafted with input from all authors.

