## Supplementary information for "PfCERLI1, a conserved rhoptry associated protein essential for invasion by *Plasmodium falciparum* merozoites"

### Supplementary methods

#### Ring-stage retention assay

To determine if successfully invaded PfCERLI1 KD merozoites developed and survived the following cycle, synchronous ring-stage parasites were set up in technical triplicate at 1% parasitaemia and 1% haematocrit either in the presence or absence of GLCN using duplicate 96-well U-bottom plates. 48 hours later, ring-stage parasitaemia was determined in one plate using flow-cytometry. In the duplicate plate, GLCN was either washed out or left unwashed and heparin was added to all wells to prevent further invasion events. 36 hours later, trophozoite-stage parasitaemia was determined by flow cytometry. To determine the percentage of rings that were retained from ring to trophozoite stages, the following calculation was used: $(\frac{Trophozoites\ (\%\ RBCs)}{Rings\ (\%\ RBCs)}) \times 100$. This calculation used the mean trophozoite and ring stage parasitaemias of the technical triplicates for each of the two biological replicates.

#### Theoretical calculation of invasion inhibition contribution to free merozoites

To calculate the build-up of free merozoites that can be detected relative to control due to invasion inhibition, the number of free merozoites was determined using flow cytometry as described above and the following equations were implemented, with the final equation applied to each experimentally determined value:

$$\frac{Parasitaemia\ fold\ change^{\#}}{Merozoites\ per\ schizont^{@}} = Prop.\,of\ produced\ merozoites\ that\ invade\ (mi)$$

$$\frac{5.5}{19.1} = 0.288$$

$$avg\ inv.inhib\ \times\ mi = Prop.\,of\ merozoites\ that\ invade\ with\ KD\ \ (msi)$$

$$0.52\ \times\ 0.288 = 0.155$$

$$mi - msi = Prop.\,of\ merozoites\ that\ fail\ to\ invade\ due\ to\ knockdown\ (mfi)$$

$$0.288 - 0.155 = 0.133$$

$$(1 - mfi) \times actual\ free\ merozoites = theoretical\ \ merozoites\ if\ no\ inv.inhib.$$

[#] as determined by [1],[@] average value from Figure 3a used.

**Prediction of PfCERLI1 signal peptide, protein structure, and ligands**

The presence of a signal peptide for both PfCERLI1 and RAP1 was predicted using SignalP-5.0 [2,3]. Full length PfCERLI1 protein sequence was submitted to the online protein structure prediction tools Phyre2 and I-TASSER. Residues corresponding to the predicted C2 and PH domains were then submitted to the online protein-ligand binding site prediction tool COACH. All predicted structures were visualised, presented and imaged using Jmol [4].

**Transmission electron microscopy**

Synchronous ring stage parasites were either treated with 2.5 mM GLCN or left untreated and incubated in standard conditions until early schizont stages before incubation with 4-[2-(4-fluorophenyl)-5-(1-methylpiperidine-4-yl)-1H-pyrrol-3-yl] pyridine (compound) 1 for 5 hours and Percoll purification of mature schizonts. Percoll purified schizonts were then fixed in 2.5 % v/v glutaraldehyde in PBS overnight at 4°C. Cells were pre-embedded in agarose and fixed in 2% osmium tetroxide reduced in 1.5% potassium ferricyanide in 0.15 M cacodylate buffer for 1 hr at room temperature. Subsequently, the cells were dehydrated in a graded series of ethanol- $H_2O$ mixtures, followed by progressive infiltration with EPON resin and embedding; 70 nm sections were prepared using an ultramicrotome (Leica EM UC7, Leica Microsystems). The sections were poststained with 4% uranyl acetate in water and Reynold's lead citrate. Thin sections were observed on a transmission electron microscope at 200 kV (Tecnai G2 F30, FEI).

**Quantitative analysis of foci and colocalisation in super-resolved PfCERLI1$^{HAGlmS}$ late-stage schizonts**

To adapt a previous workflow on the automated measurements of colocalisation in cancer cell lines [5] to malaria infected red blood cells, an advanced 3D model-based image analysis pipeline was developed in the Imaris Suite software (v9.3.0, Bitplane Inc., Switzerland) using the surface module. It accounts for significantly smaller objects, more irregular shapes and elevated levels of signal heterogeneity (e.g. 32-64 rhoptries per schizont and 0.1-0.3 µm in size depending on maturity). The approach supports quantitation of spatial positioning for a maximum four sub-cellular compartments labelled in four 3D channels (e.g. nucleus/DAPI, PfCERLI1$^{HAGlmS}$/488 nm, RAP1/594 nm and RON4/647 nm).

*Pre-processing of three-dimensional stacks*

To overcome artefacts caused by digital conversion of a light signal into an electronic signal, autofluorescence or noise, micrographs were thresholded to identify objects that are brighter than the background noise, as follows. Threshold offset was based on negative control samples (i.e. blocking peptide and/or omitted primary antibody controls) that lack cellular objects in the image. Applying a fixed threshold value [e.g. 500 for delineating RON4, 800 for RAP1 and 750 for PfCERLI1$^{HAGlmS}$ (0-32767)] allowed object detection in untreated conditions as well as in situations where images with reduced signals were expected (e.g. from a loss of label due to a compound treatment). Prior to object identification, the non-cellular background was computed and subtracted in each channel independently using a 3D surface fitting method. In this step, local background maxima around each pixel are computed and high frequency components supressed by subtracting the background from the original image. The main advantage of the method is that it minimizes the effect of the background correction (removal) procedure on the intensity values of the analysed objects. Next, identified objects were subjected to 3D Gaussian filtering (morphological) to remove very small speckles (e.g. single-voxel noise) from the image and to consolidate fragmented objects. Next, objects that touch the border of the image field were excluded from further analysis. Having identified optimal intensity thresholds and appropriate degree of smoothing for object identification, we next segmented objects in each channel to resolve and identify foci of interest as described below.

*Three-dimensional foci detection and segmentation*

For automatic detection and quantitation of PfCERLI1$^{HAGlmS}$, RAP1 and RON4 foci in the 488 nm, 594 nm and 647 nm channels, a series of 3D surface detection, 3D model-based segmentation, and surface filtering was performed using Imaris batch mode (v9.3.0, Bitplane Inc., Switzerland). Two methods of object identification were used prior to quantitative analyses: geometric (shape and size) and intensity (intensity peaks). The geometric method splits touching objects on the basis of shape, relying on boundary indentations to locate a line of separation. The intensity method separates touching objects using intensity peaks. This approach identifies each object with single, dominant intensity peak and uses the minimum relative height of the intensity peak (i.e. image contrast) for segmentation. Using this approach, objects were reconstructed as artificial 3D masks and their physical properties (e.g. intensity or morphology) adjusted to specified ranges, as follows. First, selection parameters related to object's intensity were restricted to specified ranges and those objects that are too bright or too dim were excluded from further analysis. Next, object identification was limited to foci with diameter > 0.1 µm to restrict the ranges for variation in size. Lastly, area selection parameter was used to remove noise (1-2 voxel regions). To optimise object detection and eliminate false objects, coloured solid image overlays that are exactly the size and shape of each object were used for visual inspection of segmentation accuracy and to validate gating parameters. The volume statistics exported on a per cell basis included: area ($\mu m^2$), volume ($\mu m^2$), total intensity (a.u.), sphericity (a.u.) and ellipticity (a.u.).

*Measuring colocalisation in three-dimensional stacks*

To gain a more detailed picture of how PfCERLI1$^{HAGlmS}$, RAP1 and RON4 are positioned in the rhoptries we utilised Imaris Coloc Suite (v9.3.0, Bitplane Inc.,

Switzerland). It provides an automated quantitation of colocalisation based on correlation coefficients that measure the strength of the linear relationship between two variables, i.e. the grey values of fluorescence intensity voxels of green and red image pairs. First, the two channels (e.g. 400 nm and 594 nm) for colocalisation detection are selected and voxel distribution of the two images plotted against each other as a scatter plot. The intensity of a given voxel in the green image is used as the x-coordinate of the scatter plot and the intensity of the corresponding voxel in the red image as the y-coordinate. Pearson’s correlation coefficient was then used for initial identification of diverse relationships between PfCERLI1$^{HAGlmS}$, RAP1, and RON4. It calculates the relationship between intensities in two images by linear regression. The slope of the fitted line provides the rate of association of two fluorophores, i.e. the PCC provides an estimate of the goodness of this approximation. Its value can range from +1 to -1, with 1 standing for complete positive correlation and -1 for negative correlation, with zero indicating no correlation. Scatterplots and PCC point to colocalization especially when it is complete; however, they rarely discriminate differences between partial colocalization or exclusion, especially if images are noisy (e.g. in cases where signal is dispersed or reduced due to treatment). Since evaluation of colocalization using PCC alone may be ambiguous due to variations in fluorescence intensities or heterogeneous colocalization relationships throughput the sample, we next employed Mander’s correlation coefficient (MCC) to study different stoichiometries of foci association. MCC is based on the PCC with average intensity values being taken out of the mathematical expression [6]. This coefficient varies from 0 to 1, the former corresponding to non-overlapping images and the latter reflecting 100% colocalization between both images. Because MCC is sensitive to noise, regions of interest (i.e. masking area) was defined based on fixed intensity threshold to exclude non-cellular data from further analysis. The volume statistics exported included: number of colocalised voxels (total count of colocalised voxels), % of data set colocalised (percentage of total dataset voxels colocalised), % region of interest (ROI) colocalised (Percentage colocalization of channel A and channel B volume inside the region of interest), Pearson's coefficient (PCC) in ROI volume (PCC of channel A and channel B inside the region of interest), Original Mander's coefficient (MCC) A/B and thresholded Mander's coefficient A/B.

**Supplementary Figure 1. GLCN treatment of PfCERLI1$^{HAGlmS}$ parasites specifically knocks-down PfCERLI1 expression. (a)** Synchronous PfCERLI1$^{HAGlmS}$ ring-stage parasites were either treated with 2.5 mM GLCN (+) or left untreated (-), harvested, and saponin lysed at schizont-stage in the same cycle. Parasite lysates were then used for Western blots, probed with anti-HA (PfCERLI1), anti-EXP2 (loading control), anti-GAP45, anti-RAP1 and anti-RON4 antibodies. Representative images of 3 independent experiments. **(b)** Western blot band intensities were quantified, and results are displayed as % protein expression (band intensity) in GLCN treated samples relative to untreated samples (control). (n=3, error bars = SEM).

**Supplementary Figure 2. Effect of PfCERLI1 knockdown on parasite development post-invasion and the contribution of invasion inhibition to quantified free merozoites. (a)** Quantification of free merozoites as presented in Figure 2d, but the data points in the 2.5 mM GLCN treatment represent the number of free merozoites expected after subtraction of additional merozoites that failed to invade relative to untreated controls. (n=4) (error bars = SEM). **(b)** Early PfCERLI1$^{HAGlmS/GFP}$ Ring-stage parasites were treated with glucosamine (2.5 mM GLCN) or left untreated (media). Immediately after invasion the knockdown treatment was removed (washout) or left on (no washout) and ring-stage parasitaemia was determined by flow cytometry. Schizont-stage parasitaemia was then determined 36 hours later by flow cytometry with results reported as the percentage of successfully formed rings (% rings retained) that had survived to schizont-stages (n=2) (error bars = SEM).

**Supplementary figure 3. Structural prediction of PfCERLI1 and its predicted ligands.**

Full length PfCERLI1 protein structure was predicted using Phyre2 **(a)** and I-TASSER with both prediction software's independently identifying a C2 and pleckstrin homology (PH) domain in the PfCERLI1 protein. Ribbon diagrams of the C2 domain of human itchy homolog E3 ubiquitin protein ligase (ITCH) **(b)** and PH domain of human Protein kinase B (Akt) **(c)**, which were the most similar protein structures to PfCERLI1 C2 and PH domains respectively, are superimposed (purple) onto the predicted PfCERLI1 3D structures. The C2 domain of PfCERLI1 was predicted to bind both phosphocholine (PC) and a calcium ion ($Ca^{++}$), while the PH domain was predicted to bind to inositol 1,3,4,5- tetrakisphosphate (IP4). All structures presented are ribbon rainbow diagrams, with N-terminus to C-terminus corresponding with red to blue.

**Supplementary Figure 4. Signal peptide prediction.** The presence of a putative signal peptide was predicted for (**a**) PfCERLI1 and the rhoptry luminal protein RAP1 (**b**) using SignalP-5.0.

**Supplementary Figure 5. Transmission electron microscopy of PfCERLI1$^{HAGlmS}$ schizonts.** PfCERLI1$^{HAGlmS}$ parasites were either treated with 2.5 mM GLCN or left untreated and schizonts were matured in the presence of compound 1 before fixation and analysis by transmission electron microscopy. R = rhoptry, N = nucleus.

**Supplementary Figure 6. PfCERLI1$^{HAGlmS}$ knockdown is associated with a change in the shape of the rhoptry marker RAP1.** PfCERLI1$^{HAGlmS}$ ring-stage parasites were either treated with GLCN (+ 2.5 mM GLCN) or left untreated with resulting schizonts stained with DAPI, anti-HA (PfCERLI1), anti-RAP1, or anti-RON4 antibodies. Parasites were then analysed by 3D super-resolution microscopy. **(a)** Representative images of RAP1 stained PfCERLI1$^{HAGlmS}$ schizonts. **(b)** RAP1 shape (sphericity, oblate, prolate), intensity, volume and area were quantified for GLCN treated and untreated PfCERLI1$^{HAGlmS}$ parasites (n=5). **(c)** Representative images of RON4 stained PfCERLI1$^{HAGlmS}$ schizonts. **(d)** RON4 shape (sphericity, oblate, prolate), intensity, volume and area were quantified for GLCN treated and untreated PfCERLI1$^{HAGlmS}$ parasites (n=5). Both datasets were quantified using the image feature extraction pipeline detailed in Supplementary Figure 8. (ns = $p>0.05$, * = $p<0.05$, ** = $p<0.01$, *** = $p<0.001$, **** = $p<0.0001$) (error bars = SEM).

**Supplementary Figure 7. Digital image analysis pipeline for evaluation of association frequency between two fluorescently labelled rhoptry markers. (a)** Image processing and analysis pipeline. 3D super-resolution images were captured before pre-processing to move background noise. Signals in channels of interest were then converted to objects and segmented based on minimal/maximal size, shape and intensity. Objects in different channels were then analysed to assess their overlap and colocalisation. **(b)** Using this pipeline, the number of colocalised voxels between the channels was determined. The signal of both channels was then thresholded and one of the two channels was then designated as the region of interest (ROI). The percentage colocalisation, Pearson's correlation coefficient (PCC) and Mander's correlation coefficient (MCC) inside this designated ROI was then determined. Presented data represents a comparison with PfCERLI1 as the ROI and RAP1. Each data point represents an imaged schizont.

**Supplementary Figure 8. Digital image analysis pipeline used for 3D segmentation and analysis of merozoite organelles.** Merozoite organelles imaged by fluorescence microscopy often have indistinct outlines, and image segmentation methods must be implemented to subtract background from genuine signal. **(a)** In this example, PfCERLI1 immuno-labelled with anti-HA antibodies (green) in mature schizonts have been segmented at threshold values to separate signal from noise. Signals are then converted into objects, based on minimal/maximal size, shape and signal intensity. Data can then be extracted from each of these objects, including shape (sphericity, oblate, prolate), intensity, volume, and area. **(b)** Representative super-resolution micrographs of immuno-labelled rhoptries in control (untreated) or GLCN treated (+ 2.5 mM GLCN) PfCERLI1$^{HAGlmS}$ schizonts. **(c)** Data obtained from object analyses can then be compared between two different treatments to assess the influence of the treatment on the fluorescent marker of interest (PfCERLI1).
