## Supplementary figures for "PfCERLI1, a conserved rhoptry associated protein essential for invasion by *Plasmodium falciparum* merozoites"

Supplementary Figure 1

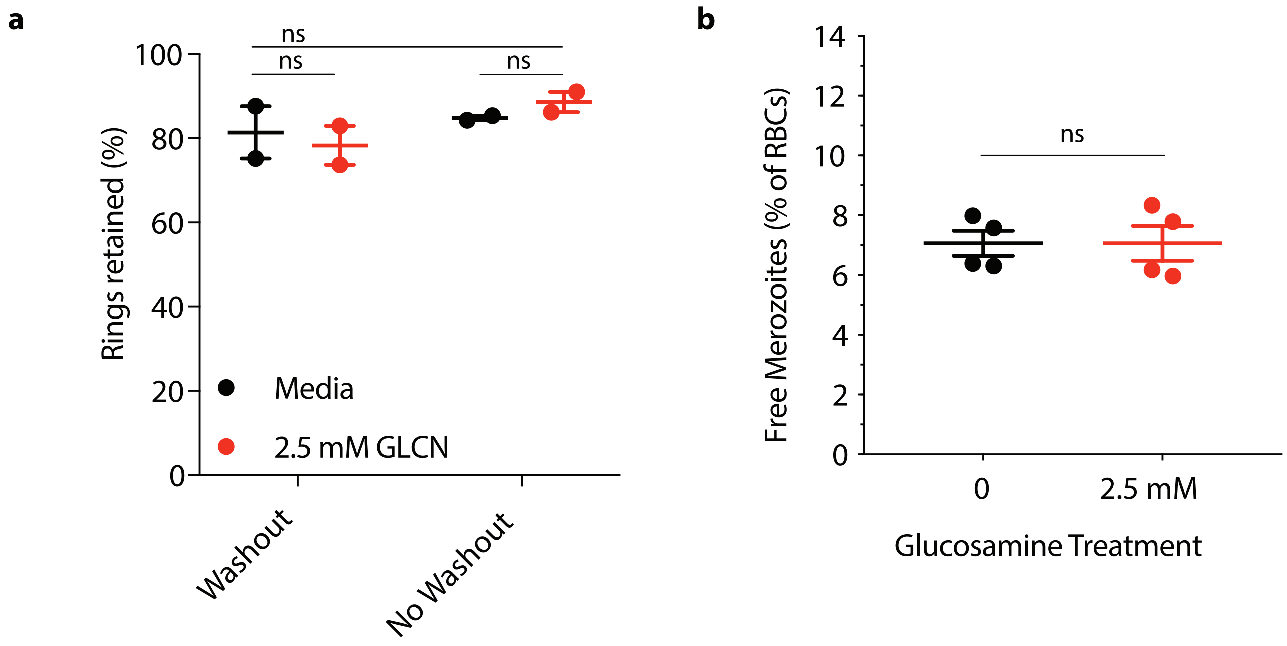

Supplementary Figure 2

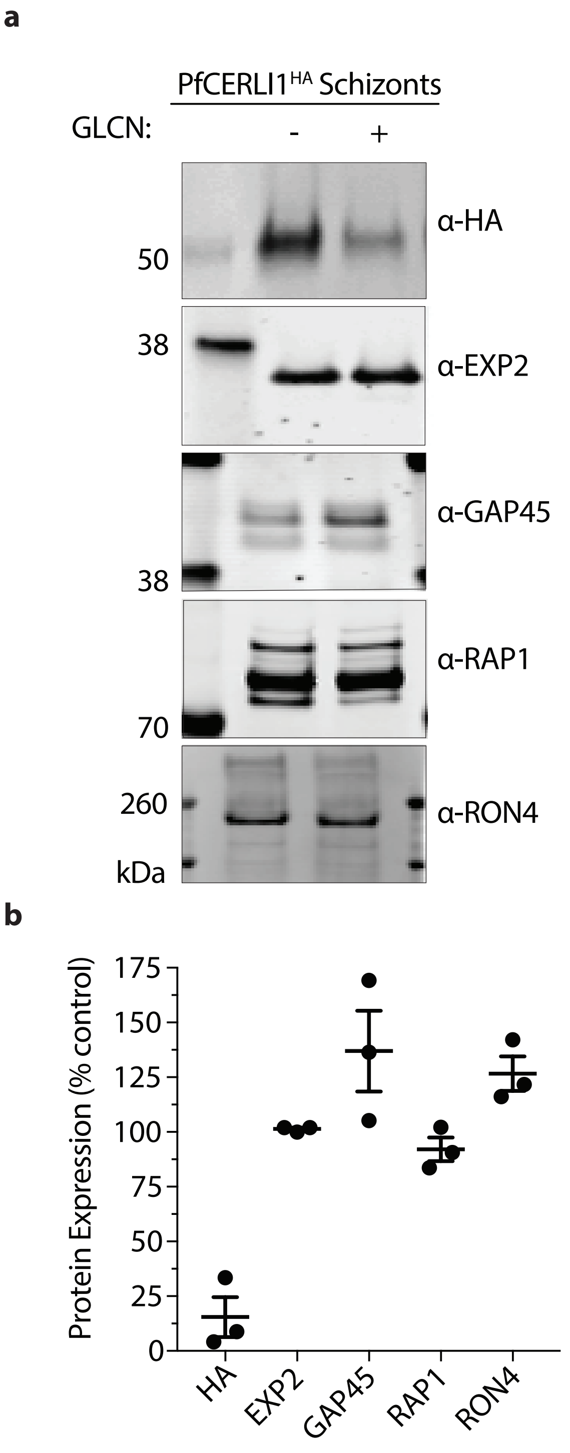

Supplementary Figure 3

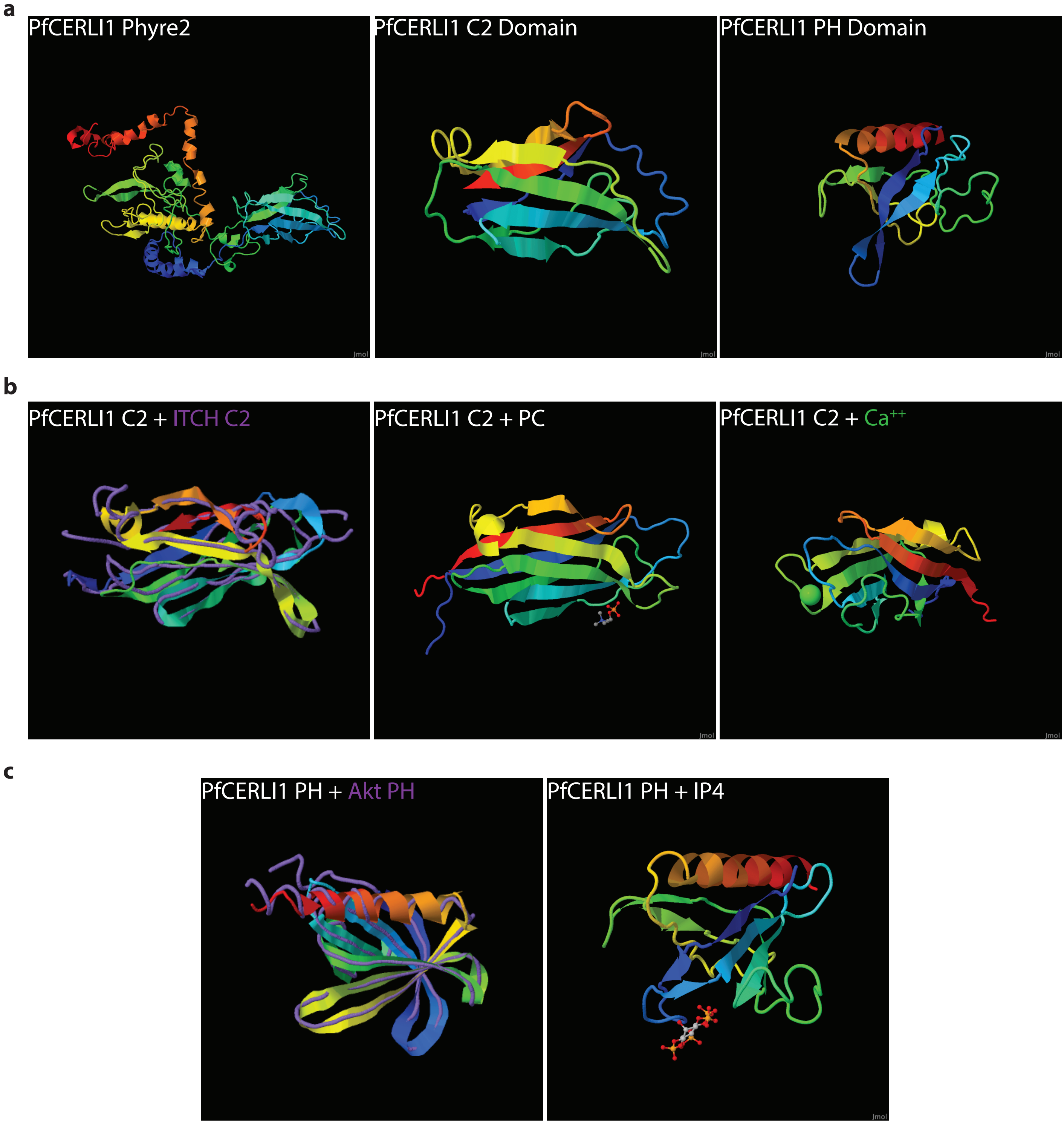

Supplementary Figure 4

**a**

SignalP-5.0 prediction (Eukarya): PfCERLI1

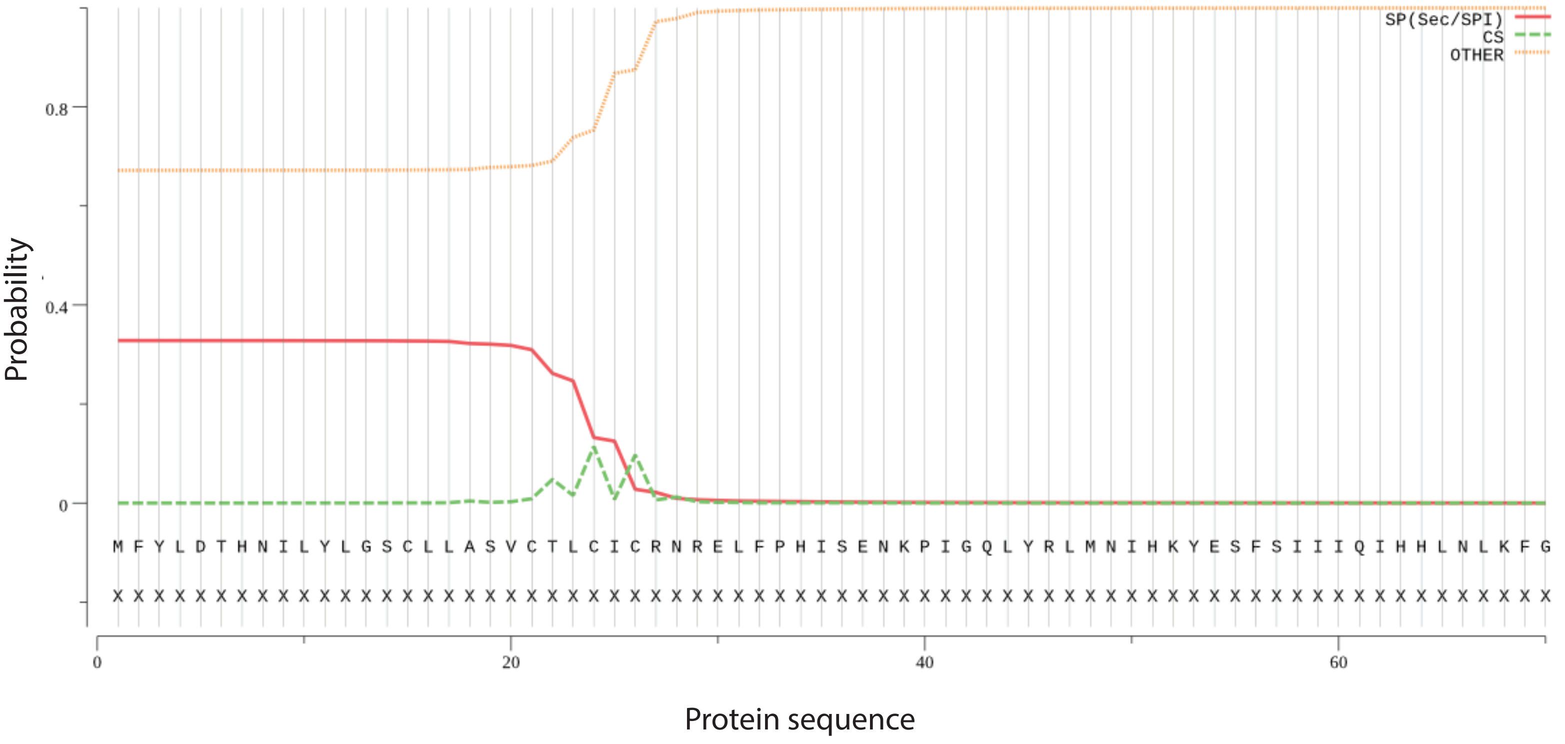


**b**

SignalP-5.0 prediction (Eukarya): PfRAP1

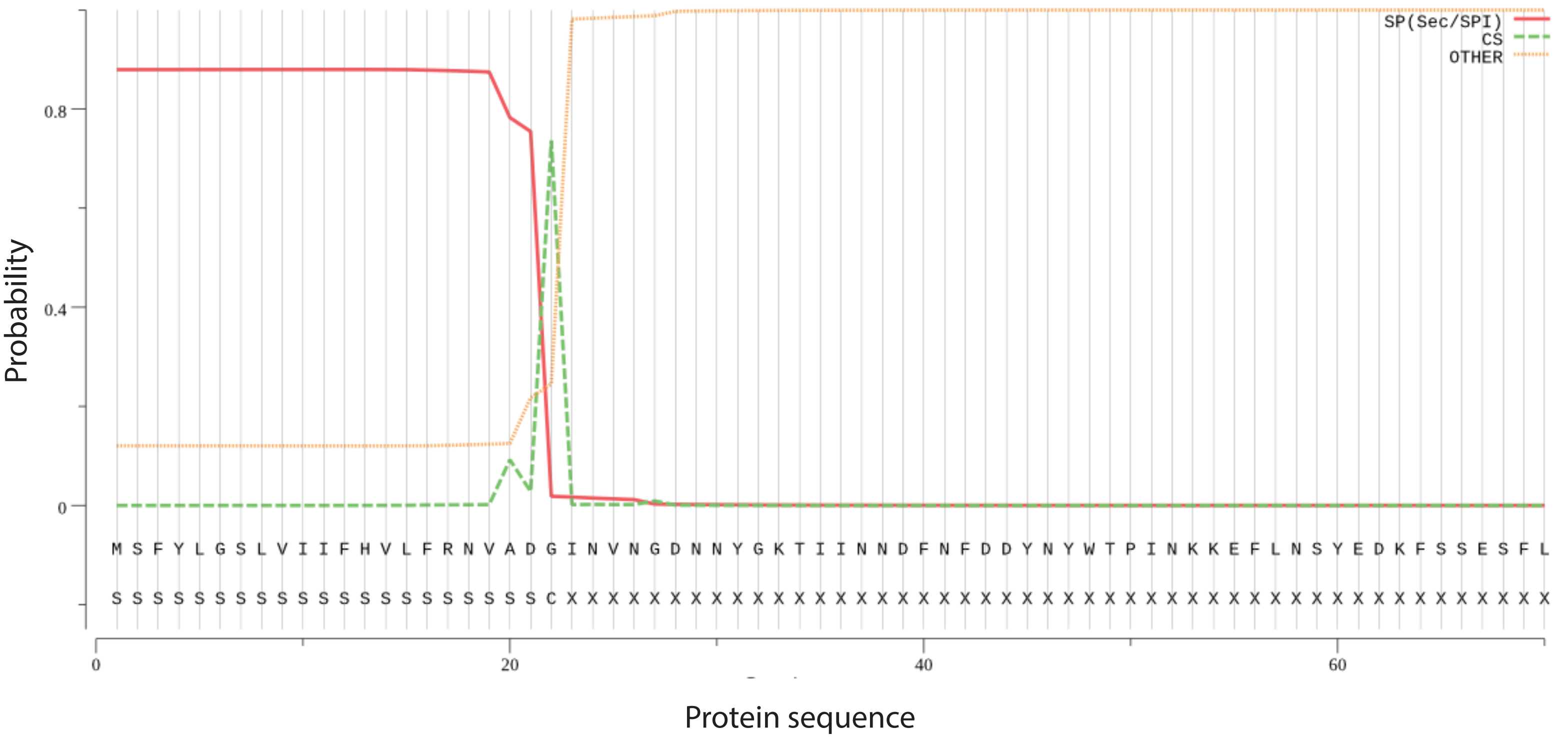

Supplementary Figure 5

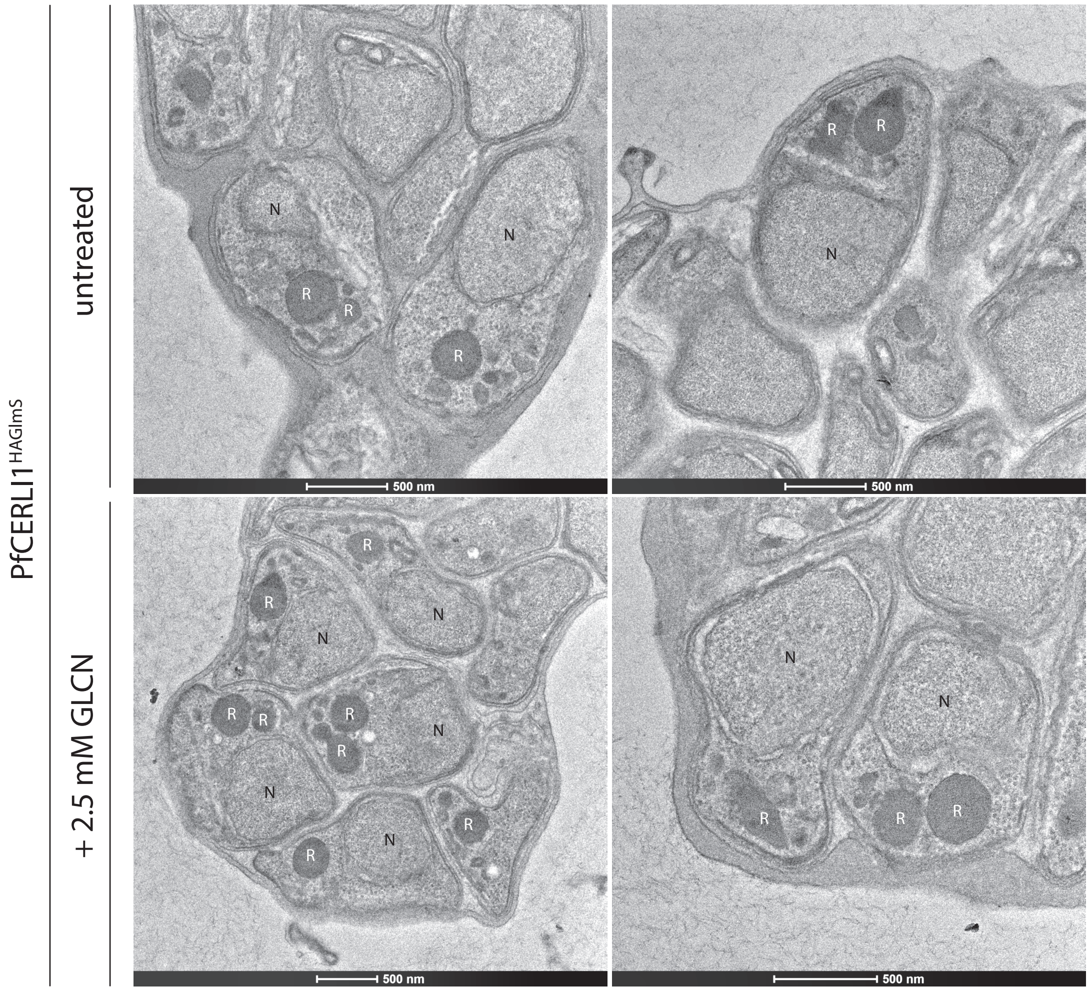

Supplementary Figure 6

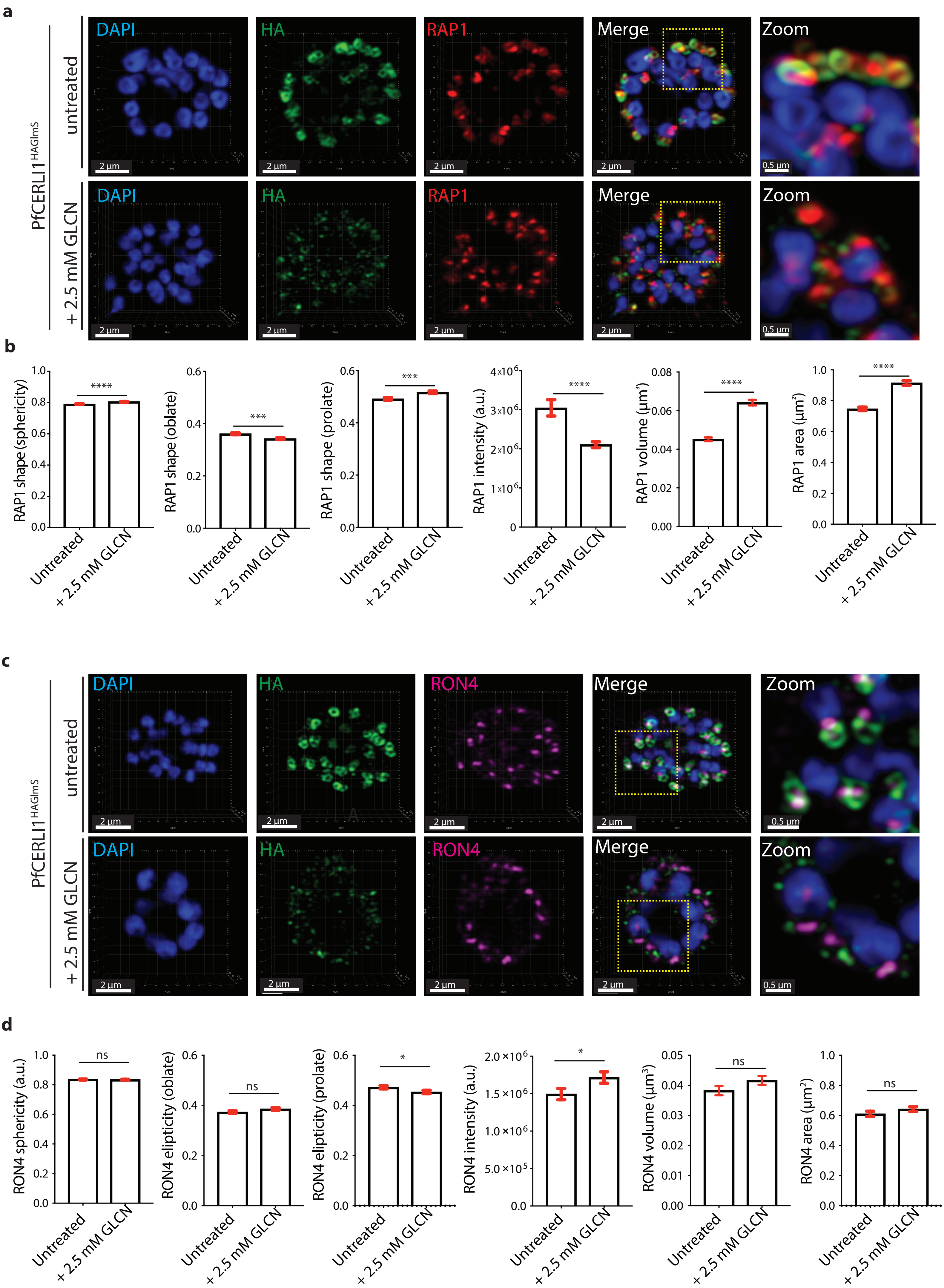

### Supplementary Figure 7

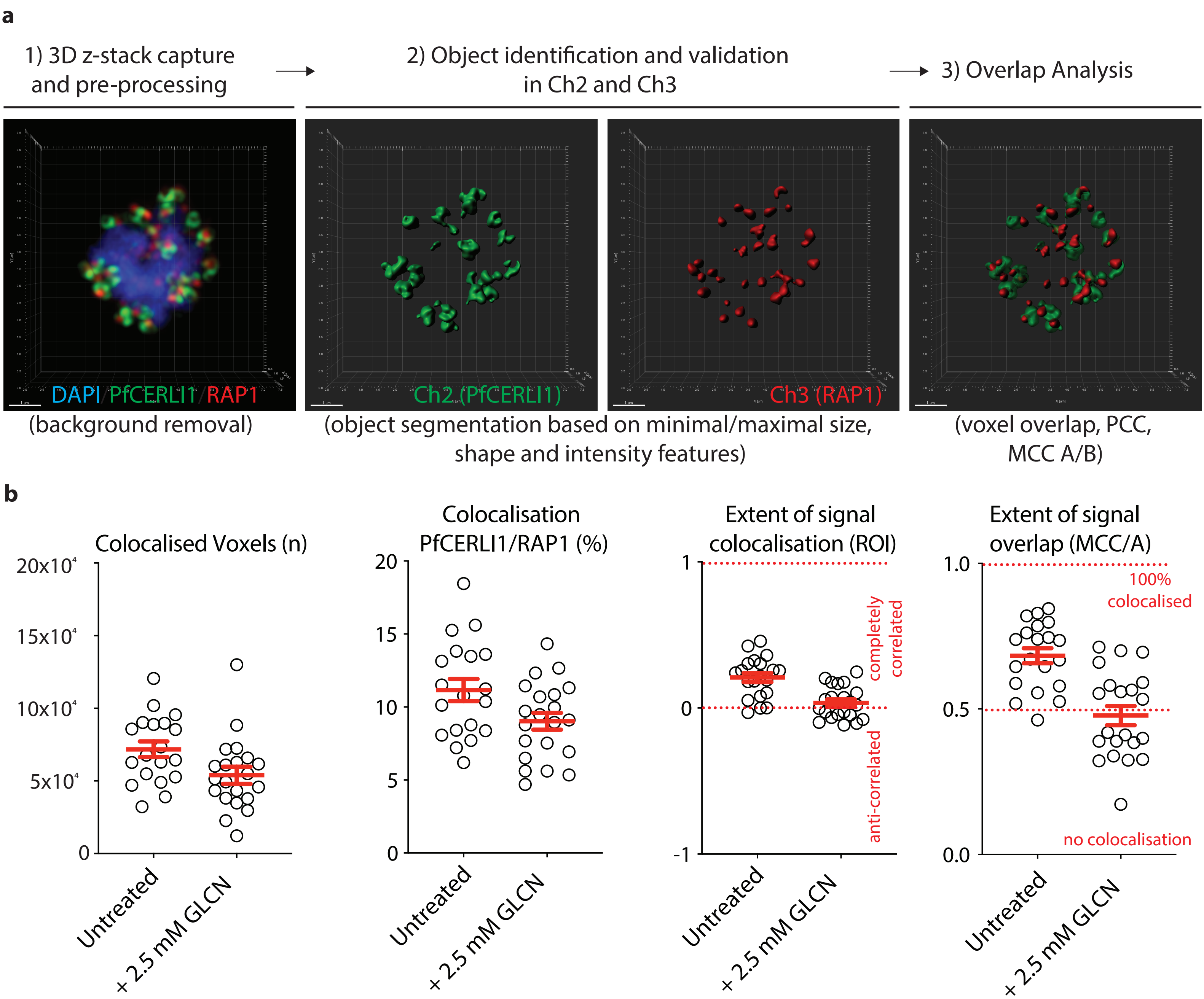

Supplementary Figure 8

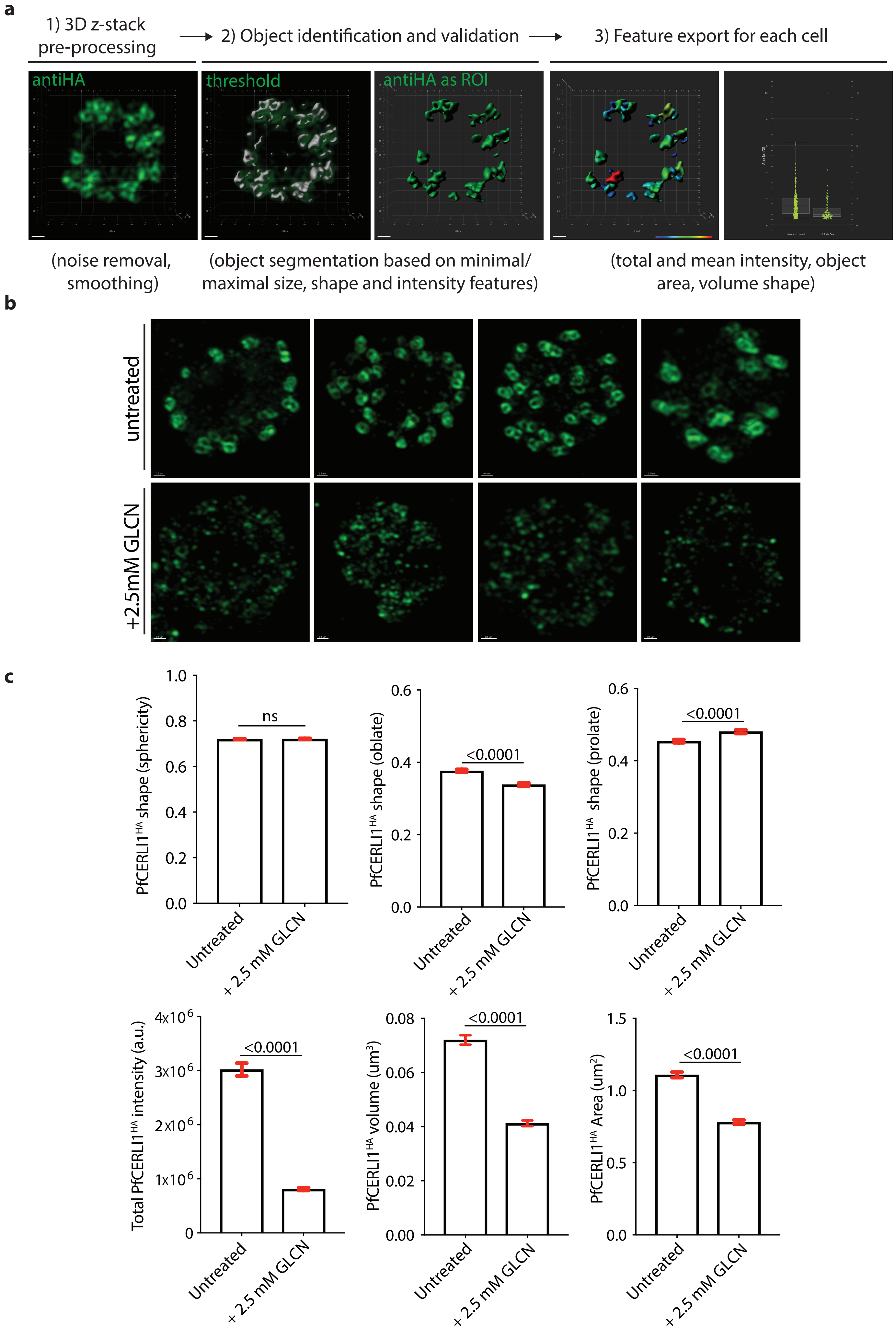
